# The amyloid concentric β-barrel hypothesis: models of Synuclein oligomers, annular protofibrils, lipoproteins, and transmembrane channels

**DOI:** 10.1101/2021.02.22.432295

**Authors:** Stewart R. Durell, H. Robert Guy

**Author notes:** Contact information: H. Robert Guy Amyloid Research Consultants (ARC), 6510 Tahawash Street, Cochiti Lake, NM 87083, USA.

## Abstract

Amyloid beta (Aβ of Alzheimer’s disease) and α-synuclein (α-Syn of Parkinson’s disease) form large fibrils. Evidence is increasing however that much smaller oligomers are more toxic and that these oligomers can form transmembrane ion channels. We have proposed previously that Aβ42 oligomers, annular protofibrils, and ion channels adopt concentric β-barrel molecular structures. Here we extend that hypothesis to the superfamily of α, β, and γ-synucleins. Our models of numerous Synuclein oligomers, annular protofibrils, tubular protofibrils, lipoproteins, and ion channels were developed to be consistent with sizes, shapes, molecular weights, and secondary structures of assemblies as determined by EM and other studies. The models have the following features: 1) all subunits have identical structures and interactions; 2) they are consistent with conventional β-barrel theory; 3) the distance between walls of adjacent β-barrels is between 0.6 and 1.2 nm; 4) hydrogen bonds, salt bridges, interactions among aromatic side-chains, burial and tight packing of hydrophobic side-chains, and aqueous solvent exposure of hydrophilic side-chains are relatively optimal; and 5) residues that are identical among distantly related homologous proteins cluster in the interior of most oligomers whereas residues that are hypervariable are exposed on protein surfaces. Atomic scale models of some assemblies were developed.

## Introduction

Amyloid-forming peptides and proteins underlie or are involved in devastating diseases including Alzheimer’s, Parkinson’s, Type 2 diabetes, and Creutzfeldt-Jakob (Mad Cow’s), and some cancers. Analyses of their structures, functions, and dysfunctions pose complicated problems that are difficult to solve. Amyloids are shape-shifters with a substantial amount of disorder. They adopt a plethora of secondary structures, conformations, and assemblies, these are often present simultaneously in dynamic equilibrium and depend upon numerous factors; e.g., peptide and salt concentrations, time, cofactors, post-translational modifications, and initial structure (seeds). α-Synuclein (α-Syn), a contributor to Parkinson’s Disease (PD), is a prime example (for recent review see Meade et al. [1]). Studies by Cremades et al. [2] illustrated the variance in oligomer sizes and how they alter as a function of time (Fig. 1). Similar results were obtained for the dissociation of fibrils into oligomers, demonstrating the dynamic nature of transitions among numerous semi-stable assemblies.

**Figure 1.**
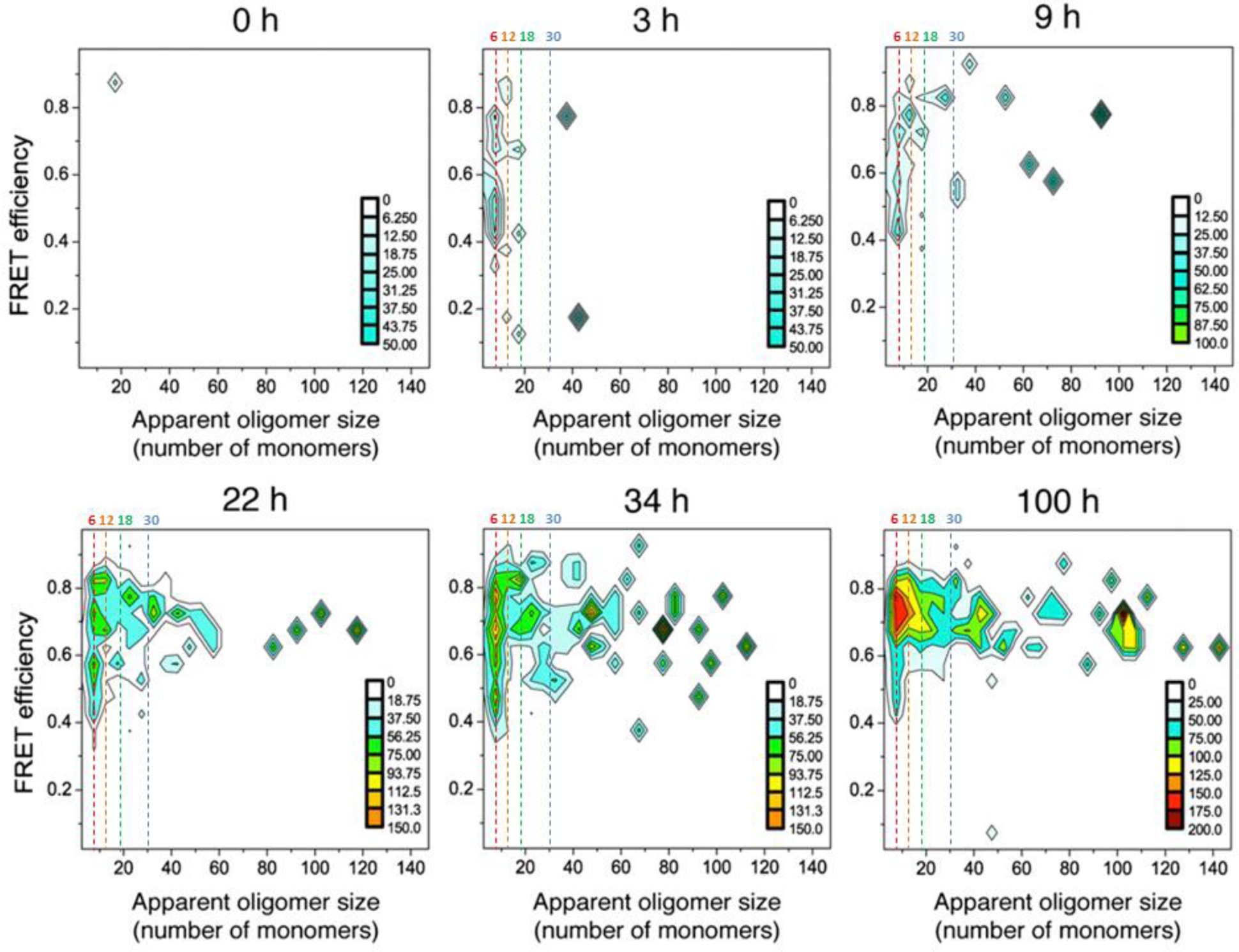
The number of α-Syn monomers/oligomer at different times during formation. Most oligomers have less than 12 monomers in studies by Cremades et al. [2] (copied with permission). We added dashed lines to indicate 6, 12, 18, and 30 monomers: the number of monomers in some of our models.

Even the secondary structures of the assemblies alter with time. Recently Zhou and Kurouski [3] used atomic force microscopy-infrared spectroscopy to analyze the structure of α-Syn oligomers at different stages of aggregation. The nanometer resolution of this approach allows different aggregate species to be analyzed simultaneously. They observed high heterogeneity. Initially most assemblies were dominated by random coil and α-helical structures, but some contained β-sheets (mostly antiparallel). With time the amount of random coil/α-helical and antiparallel β-structure decreased while that of parallel β-structures increased. By day 7 parallel β-fibrils dominated, but some spherical assemblies with significant antiparallel β-content persisted. Lashuel et al. [4] obtained electron microscopy (EM) images of α-Syn annular protofibrils that assemble to form tubular protofibrils that then morph into fibrils. Chen et al. [5] developed cryoEM images of toxic cylindrical oligomers composed of ∼ 18 and 29 α-Syn monomers. Fibrils are the only assemblies for which atomic-scale structures have been determined, and then only for portions of α-Syn [6–11]. They all include at least some segments from residues 15 – 97 that form in-register parallel cross-beta sheets. However, the exact sequential locations of the β-strands, overall folding patterns of the monomers, and interactions between fibrils vary among these structures. Also, these fibrils may not be the same as those occurring in the human brain [12]. None appear consistent with previously reported EM images of ribbons [13, 14].

But functional roles of synucleins and their pathologies may involve interactions with lipids and/or fatty acids absent in all the *in-vitro* studies mentioned above (reviewed in Mori et al. [15]). The first 95 residues of α-Syn form two antiparallel helices on the surface of negatively charged membranes [16]. These interactions are especially prevalent in presynaptic vesicles where α-Syn apparently interacts with other proteins. However, Synucleins also occur elsewhere and aggregate into larger assemblies. Lewy bodies commonly found in PD patients contain large quantities of synucleins and amyloid beta amyloids but are also chocked full of lipids, fatty acids and organelles. Meade et al. [1] observed a series of helical α-Syn polymorphs that occur in the presence of lipid vesicles. Giorgia De Franceschi et al. [17] observed similar fibrils when the ratio of docosahexaenoic acid to α-Syn is relatively low. These lipid-α-Syn fibrils have much larger diameters than those formed in the absence of such amphiphiles. The α-Syn conformation is dependent upon the ratio of lipid to α-Syn; when the ratio is high α-helical oligomers form on the surface of lipid droplets [17]. In contrast, van Diggelen et al. [18] identified two α-Syn/fatty acid oligomer assemblies that were both rich in antiparallel β-structure but had differing epitopes. Eichmann et al. [19] have published EM images of high-density lipoproteins formed by all three families of synucleins in the presence of sphingomyelin. Also, numerous studies have found that α-Syn permeabilizes membranes and forms channels in both artificial membranes [20–23], and neural cells [24–26].

Several familial mutations that contribute to PD have been identified, most of which are sequentially near each other. However, it is unclear how or whether these alter oligomer conformations or the binding of this segment into a cleft within the Cyclophilin A protein [27].

Findings that numerous α-Syn assemblies typically coexist in equilibrium, alter slowly with time, have only partially ordered and/or fractionally occupied secondary structures, and are affected by cofactors, complicates experimental determination of their structures and what makes some toxic. Also, computational modeling that relies on molecular dynamic simulations of randomized assemblies is problematic because oligomeric and fibril development often occurs over days or weeks and involves many monomers whereas most molecular dynamic simulations last less than a millisecond for a relatively small number of monomers. Furthermore, assembly conformations may depend upon factors not included in the simulations.

We utilize an alternative approach to modeling oligomers and transmembrane channels that concentrates on β-barrels, most of which are concentric. Previously we published concentric β-barrel models of amyloid-β42 (Aβ42) hexamers, dodecamers, annular protofibrils, and transmembrane channels [28–30]. Several aspects of our models including six-stranded antiparallel β-barrels and concentric β-barrels were unprecedented. Since then, however, determination of the structures of two channel-forming toxins, lysenin [31] and aerolysin [32], have demonstrated that assemblies composed of multiple identical subunits related by radial symmetry and that contain concentric β-barrels occur naturally. Also, Laganowsky et al. [33] found that a segment from an amyloid-forming protein, alphaB crystalline, can adopt a radially symmetric six-stranded antiparallel β-barrel hexamer structure (which they call cylindrin) similar to the core β-barrel of our Aβ42 hexamer model, Do et al. [34] found that the sequences of portions of Aβ can also form this motif, and Serra-Batiste et al. [35] found that Aβ42 channels have β-barrel structures with only two distinct monomer conformations.

Stimulated by these and other supportive experimental findings, we have renewed our modeling efforts. Although some details of our current generation of models have been altered and are still evolving, the basic concept of assemblies composed of concentric β-barrels [30] has been retained. Here we extend those concepts and techniques to develop models of a variety of non-fibril Synuclein assemblies. Given the difficulties described above, attempts to model these assemblies in the absence of high-resolution structural data may seem ludicrous. A saving grace is that numerous lower resolution microscopy studies have revealed circular and cylindrical assemblies composed primarily of β secondary structure. The simplest way to model circular and cylindirical β-structures is as β-barrels, which are highly constrained if all subunits are identical and they are symmetric and concentric.

## Methods, constraints, and criteria

Our analysis begins with Synuclein sequences. Fig. 2 shows our alignment of the three human Synucleins. We divided these sequences into three domains, and the domains into segments; residues 1-61 comprise the N-terminus (Nt) domain, residues 62-97 the NAC (Non-Amyloid Component) domain [1], and residues 98-140 of α-Syn comprise the C-terminus (Ct) domain. We divided the Nt and NAC domains into eight segments, Sy1-Sy8. Each segment is 11 residues long except for Sy1 and Sy5, which are four residues longer. Sy7 is missing in β-Syn, but otherwise no indels occur in the Nt and NAC domains among the three families of Synucleins.

**Figure 2.**
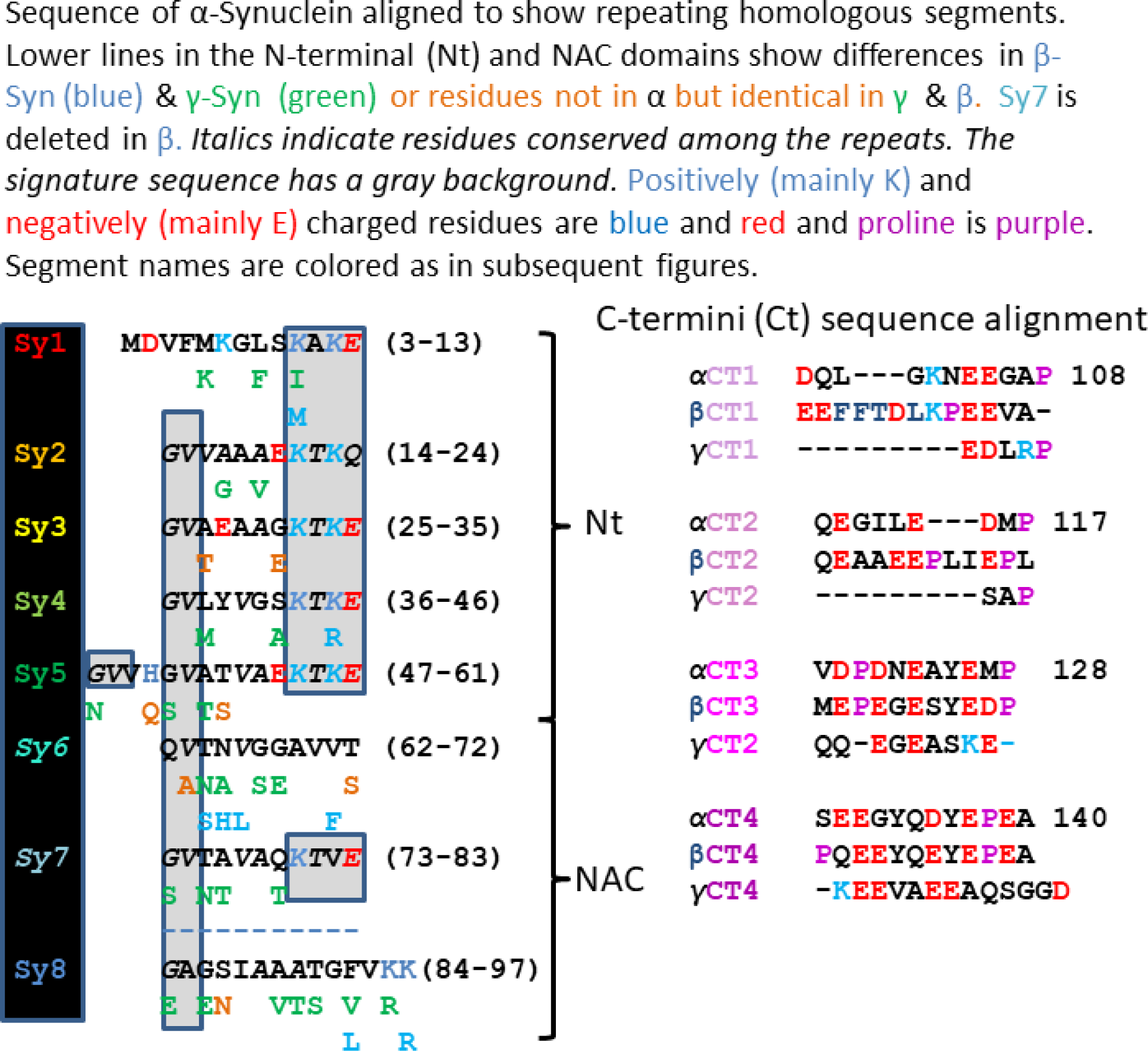
Sequence alignments of the three human Synuclein families.

Our alignment is based primarily on aligning highly conserved signature sequences (consensus KTKEGV). These are present in Sy1-5 and Sy7. Sy6 and Sy8 have some characteristics of the Nt segments but are more hydrophobic and lack the signature sequence. Ct of α-Syn appears to contain four sequentially similar segments that are nine to thirteen residues long. We call these segments Ct1, Ct2, Ct3, and Ct4 in most models. All contain an abundance of negatively charged residues (residues 104 – 140 have 15 negatively charged carboxyl groups and no positively charged groups) and a proline at or near their end. Sequences of Ct1 and Ct2 are poorly conserved between α-Syn and β-Syn with two short 3-residue insertions in β-Syn; most of the Ct1-Ct2 sequence is deleted in our alignment of γ-Syn. The Ct3 sequences are similar in α-Syn and β-Syn, and Ct4 sequences are almost identical. Ct3 and Ct4 sequences are present but not highly conserved in our alignment of γ-Syn.

Structures proposed for the segments of Fig.2 differ from one another among the numerous models presented below. The Nt domain was modeled in the following ways: (1) in αβ-barrel models Sy1-Sy3 and Sy4-Sy5 each form an α-helix, (2) in Type A (antiparallel) models Sy1 – Sy5 form a five stranded antiparallel β-sheet with the signature sequences forming turn regions, and (3) in Type P (parallel) models Sy2 and Sy4 β-strands comprise part of a β-barrel and Sy1, Sy3, and Sy5 form a parallel β-sheet that comprises part of a second surrounding β-barrel. The NAC domain was usually modeled as two β-strands that form a β-hairpin; for α-Syn the turn occurs in the polar Q79-K80-T81 region; and for β-Syn, which has no Sy7 segment, the turn occurs between Sy6 and Sy8 in the SGAGN segment which has a high propensity for turns or coils [36].

Schematic illustrations of each segment were drawn to scale: the diameter of the α-helices was 0.12 nm, the rise per residue was 0.15 nm, and the axial rotation per residue was 100⁰. For β-strands the translation per residue was 0.35 nm, the perpendicular distance between adjacent strands was 0.42 nm, and the tilt angle of the β-strands was predicted by the β-barrel theory of Murzin et al. [37]. Models were constrained in several ways: (1) All subunits were required to have identical conformations and interactions with other subunits. The resulting models have two types of symmerty. All subunits are oriented in the same direction relative to the radial axis of a core β-barrel in Type 1 models; these have M-fold radial symmetry where M is the number of monomers in the assembly. Every other subunit is oriented in the opposite direction in Type 2 models; these have M/2 radial symmetry and 2-fold perpendicular symmetry. The 2-fold axes of symmetry are located between each pair of subunits, are perpendicular to the radial axis of symmetry, and intersect the radial axis at the center of mass of the assembly; adjacent subunits are related by a 180⁰ rotation about these axes. (2)The β-barrel theory of Murzin et al. [37] was used to calculate the number of strands, N, tilt angle, α, diameter, D, and sheer number, S, of putative β barrels. Values of these parameter are limited by symmetry constraints and depend upon the number of strands/monomer in a α barrel. S/N values of the models presented here are consistent with those that have been commonaly observed; they range from 0.4 to 1.5 with the most common being 1.0 (see Table S1 of the supplement and detailed description of theory in Durell et al. [30]). The structures are even more constrained if the assembly has concentric β-barrels because all of the barrels must have the same axes of symmetry. Also, based on known structures with stacked β-sheets or concentric β-barrels, the distance of ∼0.6 nm between some closely packed β-sheets in some α-Syn fibrils (personal observation) and in some antifreeze proteins [38], and our experience in developing atomic scale models, the gap distance between the walls of the adjacent barrels should be between 0.6 and 1.2 nm. Residues with small side-chains dominate putative β-strands of Nt and NAC domins, allowing the gap distances to be less than 1.0 nm in most models. For well-known physiochemical reasons, we favor models that maximize hydrogen bonds, salt bridges, interactions among aromatic side-chains, burial and tight packing of hydrophobic side-chains, and aqueous solvent exposure of hydrophilic side-chains. Residues with polar side-chains and/or that have a high propensity for turn or coil secondary structure [36] were favored for connecting loops. We favor models in which residues that are identical among distantly related homologous proteins (*e.g.*, α-Syn and β-Syn families) cluster in the interior or at functional sites whereas residues that are hypervariable and/or where indels occur among families are located primarily on protein surfaces. We favor models in which interacting pleats of adjacent β-barrels fit between each other in a manner that reduces clashes among side-chains and in which all pleats that intersect the axes of 2-fold symmetry have the same orientation in all concentric β-barrels. In some cases β-strands were positioned and oriented to minimize stearic clashes between side-chains in adjacent β-barrels and to reduce empty cavities between hydrophobic regions of adjacent β-barrels. The final constraints are experimental: the sizes, shapes, molecular weights, and secondary structures of assemblies as determined by EM and other studies.

Atomic scale models were produced with an in-house program consistent with predicted values of S/N, α, and D and schematic models. Structures were energy minimized with the CHARMM program [39]. Molecular graphics images were produced using the UCSF Chimera package from the Resource for Biocomputing, Visualization, and Informatics at the University of California, San Francisco (supported by NIH P41 RR-01081)[40]. Image averaging using Adobe Photoshop was used on some lipoprotein assemblies.

Our β-barrel models are overly structured; *i.e.*, portions of the assemblies are likely to be less ordered *in vivo* than the models imply. Secondary structures of some regions may be fractionally occupied; too dynamic for their atomic structures to be resolved, but sufficiently ordered to contribute to the overall structure and electrostatic interactions of the assembly. Also, when constraining models to correspond to sizes and shapes observed in EM studies it seems prudent to include all of the protein sequence, even if the outer portions of the models have greater uncertainty, disorder, and only approximate the general location of the region.

### Similarities to amyloid-β models and sequences

In noting the 11-residue repeating segments of Synucleins (segments Nt and NAC), authors often point to the similarity to apolipoprotein sequences. This is understandable because portions of apolipoprotein sequences do have some 11-residue amphipathic α-helical repeating segments and α-Syn adopts a conformation with two antiparallel α-helices upon interactions with negatively charged membrane surfaces [16]. Furthermore, in early stages of development α-Syn can form small oligomers that fractionally occupy helical secondary structures [41–43]. These dynamic oligomers gradually morph into primarily β oligomers, possibly transitioning through a 310 helical phase [41, 44]. Helical tetramers have been touted to be the primary native form in human brains [41, 44], but others claim it is disordered monomers [45]. In general, the presence of α-helices in α-Syn oligomers appears to dissipate with time as β secondary structure increases.

Although the positions of charged residues that make the helices amphipathic are conserved among the α-Syn and β-Syn families, the following aspects indicate that much of the sequence conservation has occurred to preserve other types of structures: 1) Nt has ten valines and only two leucines even though leucine is more hydrophobic and has a higher propensity for α secondary structures whereas valine has a higher propensity for β secondary structure [36]. 2) The threonine of the signature sequences occur on the hydrophobic face of the α-helices even though it has a higher propensity for β-secondary structure and has a polar hydroxyl group. 3) All signature sequences contain a glycine even though glycine has a much higher propensity for coils and turns than for α secondary structure. 4) Six conserved hydrophobic side chains occur on the polar faces of the Nt α-helices.

Similarities between Synuclein and Aβ sequences generally have been overlooked. In modeling of Aβ42 assemblies we divided the sequence into three segments of about the same length (Fig. 3b). The second, Aβ2, and third, Aβ3, of these segments form β-strands in numerous fibril structures [46–48], and the first, Aβ1, has a bent β structure in some fibrils [49, 50]. In our models of Aβ42 hexamers, Aβ3 forms a hydrophobic six-stranded antiparallel β-barrel that is surrounded by an antiparallel β-barrel formed by Aβ1 and Aβ2 (Fig. 3a). The 3-fold radial symmetry and 2-fold perpendicular symmetry of this Type 2 model allows all subunits to have the same conformation and interactions. Sy2 and Sy4 sequences of α-Syn are quite similar to those of Aβ2; *i.e*., they have a 5-6 residue-long hydrophobic segment (depending upon how one classifies glycine) flanked by charged residues (Fig. 3b). Sy1, Sy3, and Sy5 segments have a similar sequence except for a charged residue in the middle of the hydrophobic region. This gives them a slight resemblance to Aβ1. Two portions of the sequence of the NAC domain have longer hydrophobic regions that are quite similar to that of Aβ3; the three sequences are dominated by glycine, alanine and residues with moderate-sized side-chains that have high β propensity because they branch at the β-carbon (threonine, valine, and isoleucine).

**Figure 3.**
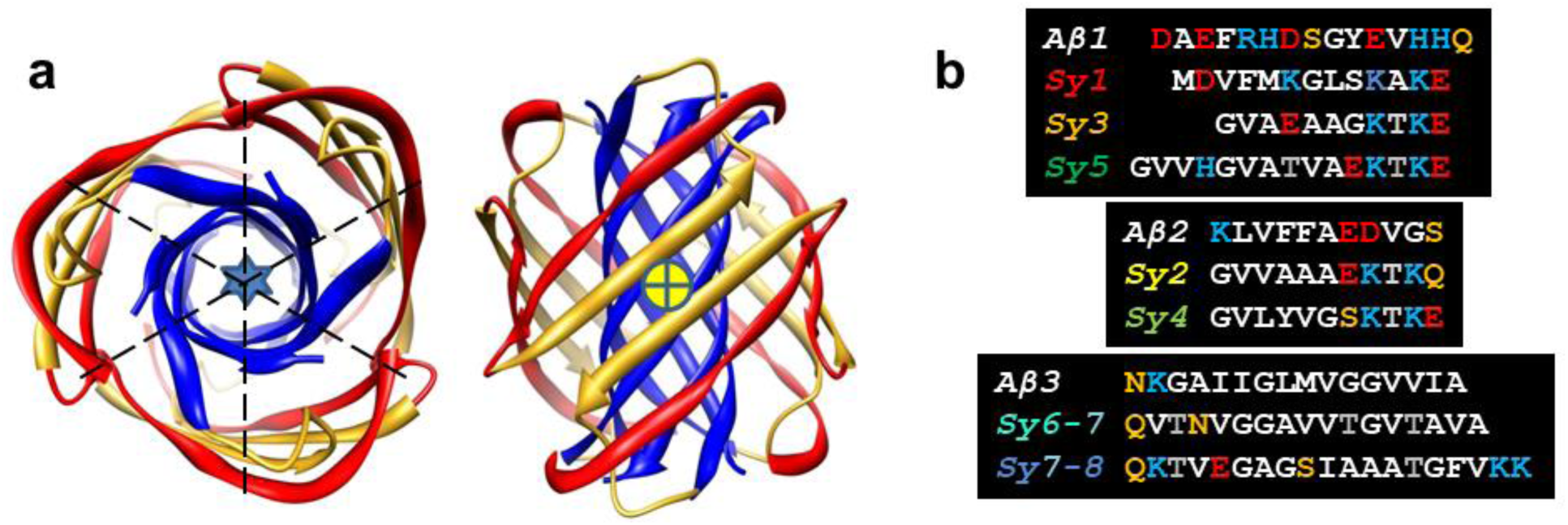
(a) Our 2010 model of a Type 2A Aβ42 hexamer; the Aβ1 strand is gold, Aβ2 is red, Aβ3 is blue, the blue star shows the location of the radial axis, the dashed lines and the + in the yellow circle indicates the locations of the perpendicular axes. In some alternative models the Aβ1 segment forms a β-hairpin. (b) Alignment of three Aβ42 segments with α-Syn Nt and NAC segments; all residues of both sequences are included. Residue color code: negatively charged, positively charged, uncharged polar, threonine, apolar and hydrophobic.

## Results

### Small Synuclein Oligomers

Some medium to large α-Syn oligomers appear to contain aggregates of smaller components [51][52, 53]. The smallest of these components (other than monomers) appear as beads of ∼ 4 nm in diameter that cluster or string together (Fig. 4). Spheres of this size could contain Nt and NAC domains of three or four monomers. The oligomers also appear to contain less defined regions surrounding the beads. These regions may contain less ordered portions of α-Syn oligomers, e.g., Ct domains, and disordered monomers; if so these disordered regions would contribute to experimental findings that the oligomers have a higher content of disordered structure than present within our models of the beads.

**Figure 4.**
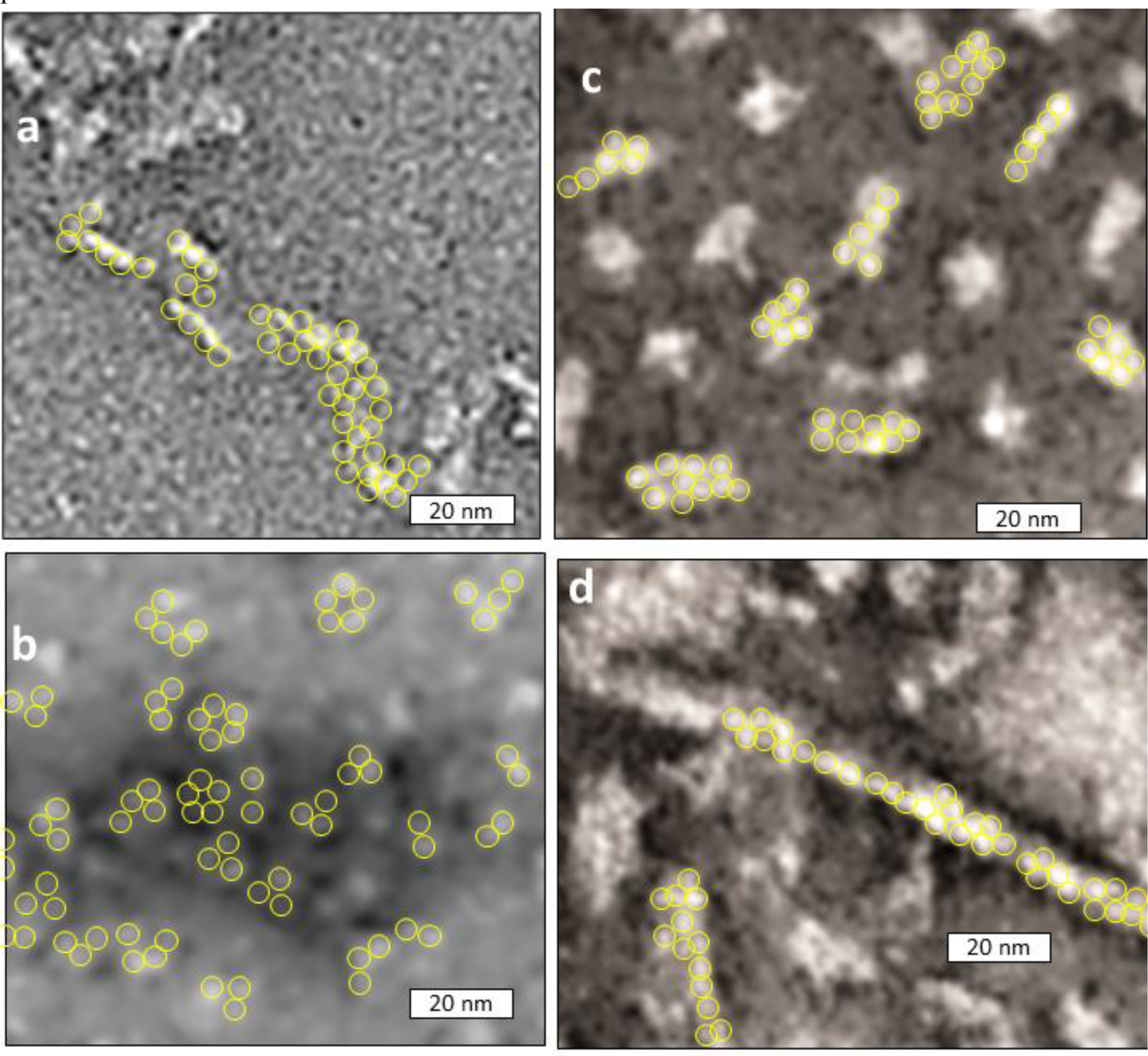
EM images of α-Syn oligomers that appear to contain circular or spherical oligomers with a diameter of ∼4 nm. Portions of published images were enlarged and adjusted to have the same scale. We added yellow circles of 4.0 nm diameter around some beads. (a) An assembly in which oligomers were stabilized using formaldehyde cross-linking [51]. (b) Adapted with permission from Paslawski W, Andreasen M, Nielsen SB, et al. High stability and cooperative unfolding of alpha-synuclein oligomers. Biochemistry. 2014 Oct 7;53(39):6252-63 [52] Copyright 2014, American Chemical Society. (d-e) small and large oligomers reported by Lorenzen et al. [53] Adapted with permission from Lorenzen N, Nielsen SB, Buell AK, et al. The role of stable alpha-synuclein oligomers in the molecular events underlying amyloid formation. J Am Chem Soc. 2014 Mar 12;136(10):3859-68. Copyright 2014, American Chemical Society.

### TYPE 2 Models

#### α-helical and α-β-barrel tetramers

Fig. 5a shows an EM image of an α-Syn tetramer composed primarily of partially occupied α-helices [41]. This image is shown to emphasize the importance of tetramers and to suggest that helical structures may represent a starting point from which oligomers with β-secondary structures develop. This type of conformational change may initiate transitions from predominately α-helical monomers with intra-monomer backbone H-bonds to parallel β-sheets of fibrils with inter-monomer backbone H-bonds.

**Figure 5.**
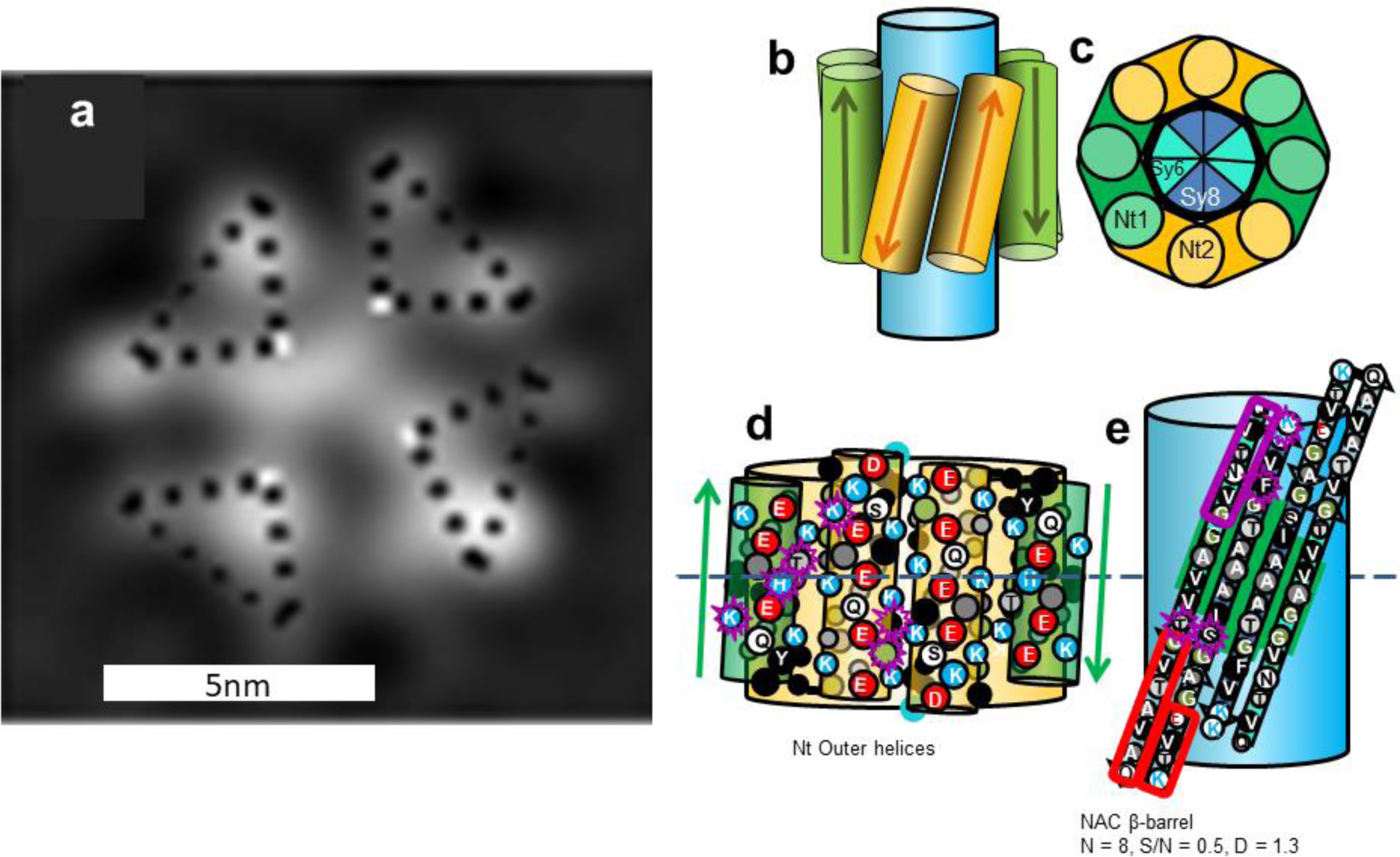
(a) EM images of fractionally occupied helical tetramer with 2- or 4-fold symmetry, Copied with permission from Wang et al. [41], and (b-e) schematics of an αβ-barrel Type 2 tetramer model for α-Syn. (b) The model viewed from the side; the large blue cylinder represents the NAC β-barrel. (c) Wedge representation of a cross-section of four Nt helical hairpins surrounding an eight-stranded antiparallel β-barrel formed by four NAC β-hairpins. (d) Representation of Nt helices for two subunits superimposed on the Nt helical-barrel. The dashed line is the plane of 2-fold symmetry. (e) Flattened representation of two NAC β-hairpins superimposed on a light blue β-barrel. The Sy7 hairpin (boxed in red) is not present in β-Syn, but it is preceded and followed by residues that are identical in α- and β-Syn (green background). The larger circles represent side-chains oriented outwardly on the front side of the helices or strands, the smaller circles are for those on the back side. Side-chains are colored by their properties: hydrophobic side-chains (V, I, L, M,F, Y) are black with white letters, less hydrophobic alanine is dark gray with white letter, glycine is green with white letter, ambivalent threonine is light gray with black letter, uncharged hydrophilic S, Q, and N, are white with black letter, negatively charged D and E are white with red letters, positively charged K and R are white with blue letters, sometimes positively charged H is light gray with blue letter. The N and C terminal residues are outlined in blue and red to indicate their positive and negative charges. Residues that differ in α- and β-Syn are encircled in purple for the subunits on the left. The number of β-strands, N; sheer number to N ratio, S/N; and diameter of backbone, D, are listed below the NAC β-barrel.

The transition from primarily α-helical to antiparallel β oligomers may involve oligomers with both types of secondary structures. A relatively common protein structural motif is a TIM barrel in which the secondary structure alternates between α-helices and β-strands, with the β-strands forming a relatively hydrophobic 8-stranded parallel core β-barrel that is surrounded by eight relatively amphipathic α-helices [54]. The model we suggest here differs in that the Nt domain forms an α-helical hairpin and the NAC domain forms a β-hairpin. Once four monomers aggregate to form tetramers, the more hydrophobic NAC domains may migrate to the core where they form an 8-stranded antiparallel β-barrel that is surrounded by amphipathic α-helices. The charged side-chains of the Nt helices would be oriented outwardly where they are exposed to the aqueous phase and can interact with oppositely charged side-chains of adjacent α-helices, and their hydrophobic surfaces on the opposite side of the helices would interact with the NAC β-barrel (Fig. 5 b-e). The α-Syn and β-Syn proteins diverged long ago [55]; nonetheless, in the Nt domain only six side-chains differ between the two families; these are all on the polar faces of the helices (circled in purple in Fig. 5). The NAC domain is less conserved; the first five residues of Sy6 differ, the Sy7 (boxed in red) putative β-hairpin is deleted in β-Syn, and three residues of Sy8 differ. However, there are stretches of four (GAVV) and six (IAAATG) consecutive identical small apolar or ambivalent (T) residues in Sy6 and Sy8 that are spatially adjacent at the center of the model (green background in Fig. 5d): alanines comprise the central pleat and alanine, glycine, and valine comprise the two adjacent pleats. Some of these conserved apolar residues should interact with conserved apolar residues on the back sides to the Nt helices. The γ-Syn sequence has more differences, but the overall conservation pattern is similar. There are 13 differences in the Nt domain, but only two are on the back side of the helices. Only 16 of 34 residues in the NAC domain are identical in the two sequences, but three and two consecutive residues are identical in the mid regions of Sy6 and Sy8.

### Concentric β-barrel Models

The general monomer backbone folding motifs used in most of our concentric β-barrel models of synucleins are illustrated in Fig. 6. These were designed so that most of the repeating segments have similar backbone conformations. The initial, relatively hydrophobic portions of the Nt segments form β-strands, and the more polar signature sequences at the ends connect these strands. Sy7 forms a β-hairpin in most cases and its two halves are often extensions of the S6 and/or S8 β-strands.

**Figure 6.**
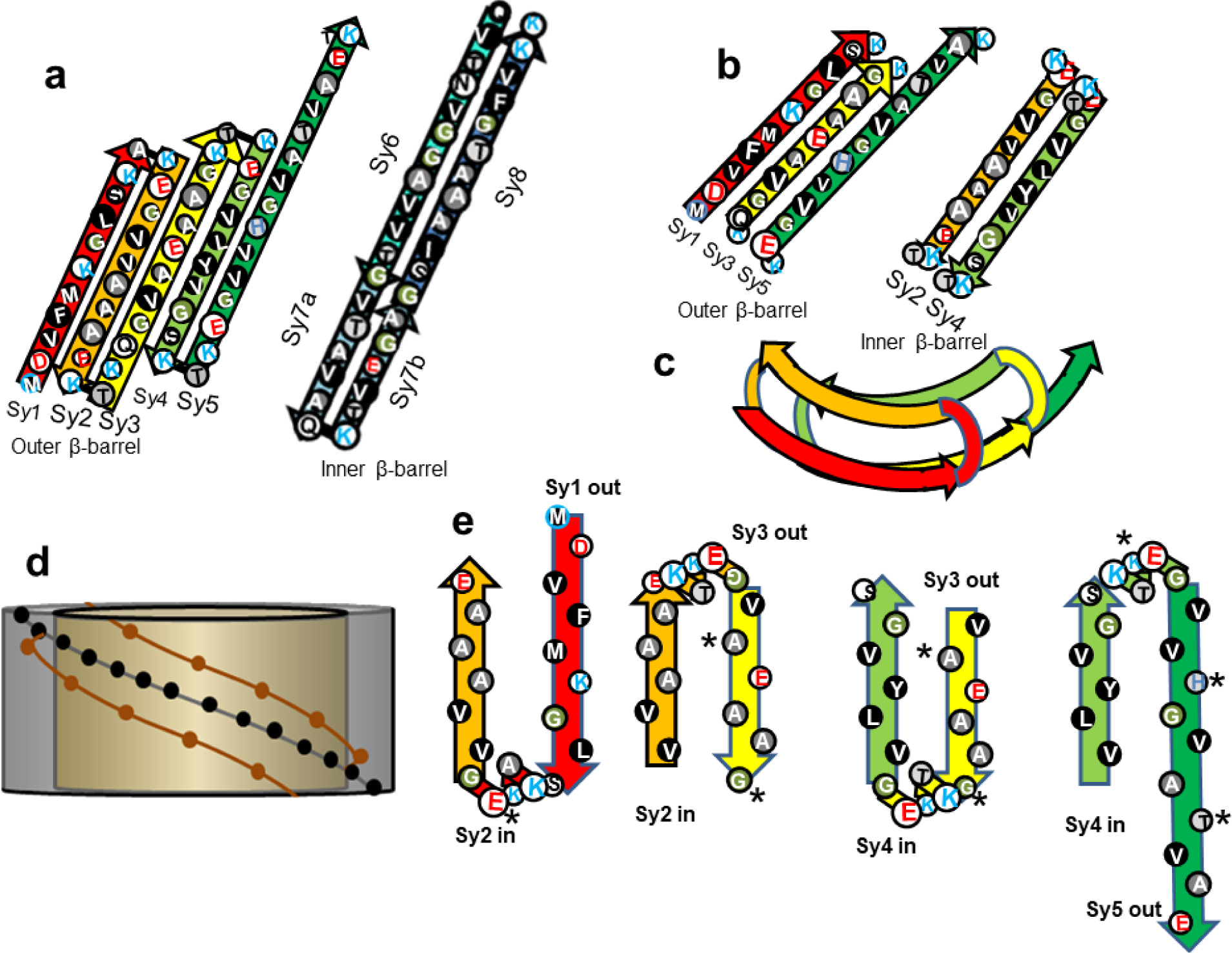
Schematic representations of a proposed structure for the Nt domain of Type A and Type P models. (a) A flattened representation of the β-strands for one subunit’s contribution to two Type A concentric β-barrels as viewed from outside the barrels. In some models S5 is part of the inner barrel and Sy7 is not always an extension of Sy6 and Sy8. (b) A flattened representation of the β-strands for one subunit’s contribution to two Type P concentric β-barrels. Only Nt segments are shown because relative positions of NAC strands vary among models. (c) Schematic of the conformation of a monomer in putative concentric Type P β-barrels as viewed down the central axis of the barrels. The strands within each barrel are parallel and oriented in opposite directions in the adjacent barrels. (d) Illustration of intermeshing pleats for concentric β-barrels that have pleats with similar pitches. The gold cylinder represents an inner barrel with the brown circles representing outwardly oriented side-chains of two adjacent pleats. The gray cylinder represents a surrounding barrel with the black dots representing side-chains of a central pleat that fits between the two pleats of the inner barrel. (e) Side view of sequentially adjacent strands approximating how their side-chains may pack between one another in Type P models. The asterisks indicate where the sequence of β-syn differs from that of α-Syn.

In most of our Type A models, the Nt domain forms a five-stranded antiparallel β-sheet; these assemble into an antiparallel β-barrel that usually surrounds a central hydrophobic core antiparallel barrel formed by the NAC domains (Fig. 6a). In Type P models Nt segments form portions of two parallel (except where subunits interact in Type 2 models) β-barrels with Sy2 and Sy4 in an inner barrel, and Sy1, Sy3, and Sy5 in the surrounding barrel (Fig.6 b,c,e). Except for small differences in strand tilt and curvature, the Nt domain folding motif is the same for all Type P models that follow. Sequentially adjacent strands are located in different β-barrels and connected by the most highly conserved and charged segments among the repeats, the signature linkers. Locations of NAC strands vary among the models.

Interacting side-chains of sequentially Type P adjacent strands may fit between one another (Fig. 6e) in a manner similar, but not identical, to steric zippers [56]. The major differences are that the interaction between side-chains of adjacent β-barrels is not one-to-one and the differences in tilts of the strands can affect side-chain packing. The pitch of a β-barrel’s pleats as they spiral around the barrel can be approximated by P = C tan α where P is the pitch, C is the circumference of the barrel, and α is the tilt angle of the strand determined by the S/N ratio, where S is the sheer number of the barrel and N is its number of strands. When P is about the same for adjacent concentric barrels, the pleats can fit between each other similar to the threads of a bolt fitting into a nut (Fig. 6d); we refer to this situation as intermeshing pleats. This is more probable when α of the surrounding barrel is less than that of the more interior barrel. In Type 2 models some pleats intersect an axis of 2-fold-perpendicular symmetry present in all the barrels. For the pleats of concentric barrels to intermesh at these points the pleats of all concentric barrels must have the same orientation (inward or outward) at these regions. This criteria places additional constraints on S/N values of the concentric barrels. The small size of the putative interacting side-chains in the Syn models likely facilitates tight packing between the barrels and reduces steric clashes among side-chains, especially when P values of adjacent β-barrels differ substantially.

### Type 2A Models

#### tetramers

The Type 2A tetramer model of Fig. 7 resembles the α-β-barrel of Fig. 4 except that the Nt domains have transitioned from helical hairpins to 5-stranded antiparallel β-sheets. The NAC β-hairpins still form an 8-stranded antiparallel core β-barrel, but now it is surrounded by a 20-stranded antiparallel β-barrel formed by Nt. Most side-chains of the central interior of the inner NAC barrel are small and apolar, alanines and glycines. The interacting faces of the pleats are composed primarily of small side-chains (G, A, V, and T for the inner barrel and G, A, V, and S for the outer barrel), facilitating relative tight packing between the barrels (gap distance ∼ 0.95 nm). The major advantages of this model are its simplicity, the facts that the most hydrophobic segments form its core and most polar side-chains are on the surfaces, the negatively charged glutamate side-chains of the signature loops can all salt-bridge to positively charged lysines, and the core NAC β-barrel is the same as proposed for the α-β-barrel tetramer model.

**Figure 7.**
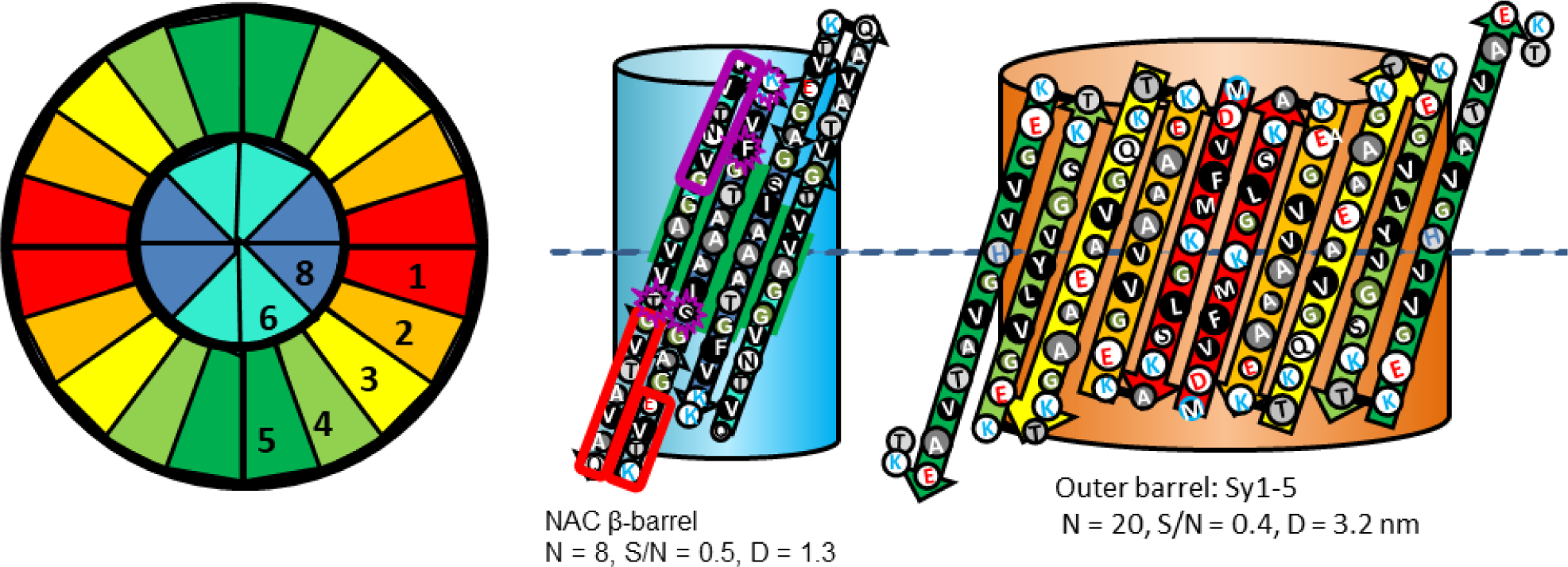
Type 2A tetramer α-Syn model. Wedge cross-section illustrates the relative positions of the β-strands. The flattened schematics represent two subunits on the front of the assemblies superimbosed on barrels. A relatively long hydrophobic 8-stranded antiparallel β-barrel (same as in Fig. 4) is formed by NAC β-hairpins at the core and an outer 20-stranded antiparallel β-barrel is formed by Nt. The dashed line in the flattened models indicates the plane of 2-fold symmetry.

#### Models of α-Syn with fatty acids

Fatty acids facilitate formation of α-Syn oligomers [57] but not β-Syn oligomers [58]. van Diggelen et al. [18] have isolated two conformationally distinct α-Syn oligomers: one complexed with the naturally occurring polyunsaturated fatty acid docosahexaenoic acid (DHA) and the other with the lipid peroxidation product 4-hydroxynonenal (HNE). The sequence-independent polyclonal A11 antibody recognizes numerous soluble amyloids, possibly by recognizing antiparallel β-sheets. It recognizes HNE-αSOs (α-Syn oligomers), but not DHA-αSOs. They report that both types of oligomers have a relatively high content of β-structure; DHA-αSOs have 62.3% random coil, 12.8% α-helical and 12.7% β-sheet and HNE-αSOs have 45% random coil, 14.6% α-helical and 28.4% β-sheet. Circular Dichroism and FTIR spectra analyses indicate the presence of antiparallel β-structure in both types of oligomers. Transmission Electron Microscopy images (Fig. 8 g & h) suggest that both types of oligomers are spherical with average diameters of 20.0±4.6 nm for DHA-αSOs and 19.5±6.3 nm for HNE-αSOs. However, these images also have much smaller bright spots that appear to self-associate. The circles in Fig. 8 indicate that the spacings between these spots are consistent with models of interactions between the Nt-NAC beta barrels with diameters of 6.0 nm for Type 2A hexamers (red) for HNE-αSOs and 7.1 nm for Type 2A octamers (yellow) for DHA-αSOs. Like EM images of numerous other oligomers (see Fig. 4), strings or clusters of apparently dense particles are surrounded by other material; possibly disordered Ct domains and entrapped monomers and/or fatty acids that may compose the remaining portions of the 20 nm diameter spheres.

**Figure 8.**
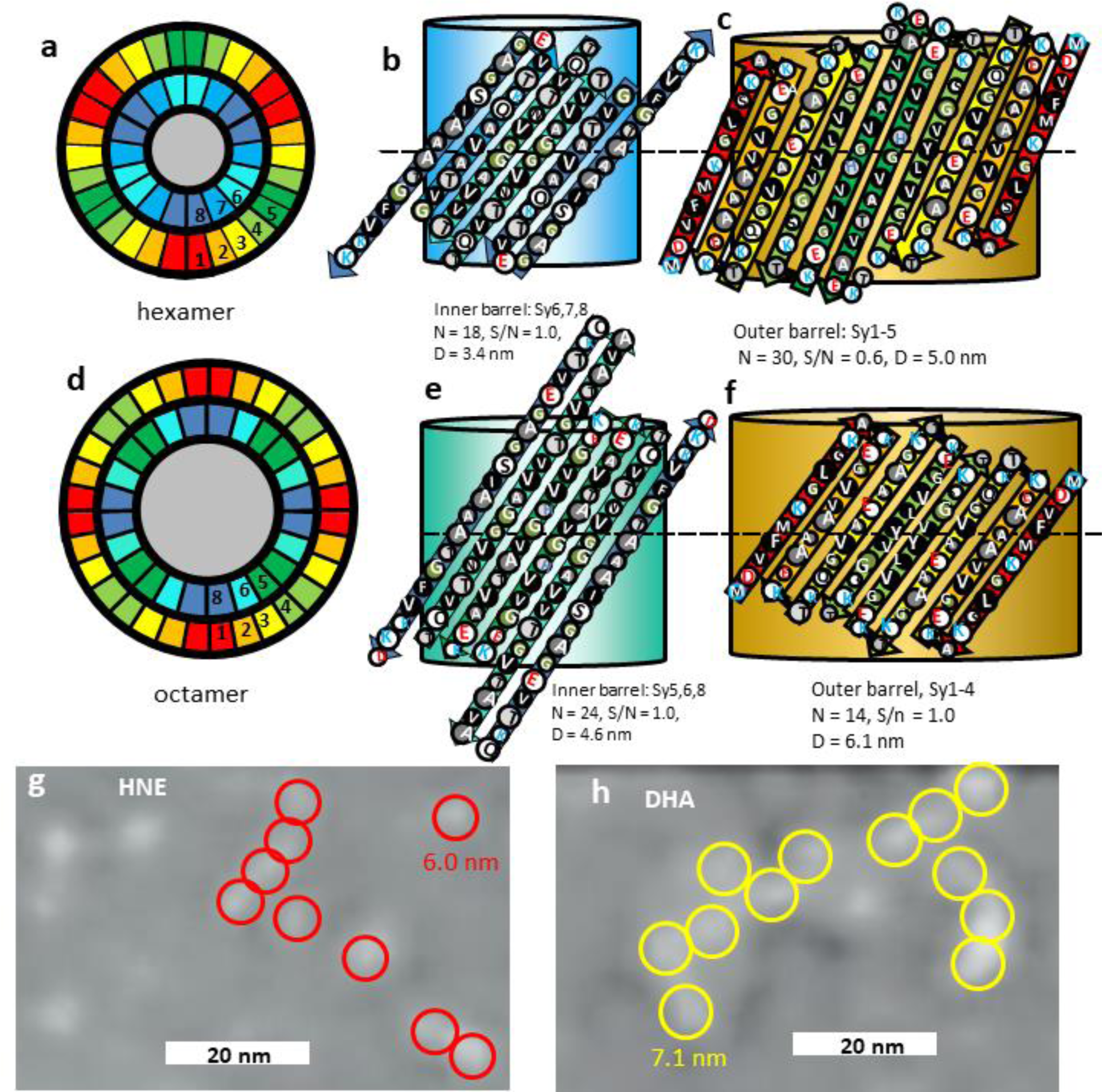
Type 2A models of hexamers and octamers and EM images of α-Syn-fatty-acid complexes. (a-c) Hexamer model. (a) Wedge representations showing the relative positions of β-strands in a cross-section. The gray region in the center represents fatty-acid alkyl chains. (b & c) Flattened representations of two subunits of the inner barrel formed by Sy6,7,8 strands and outer barrel formed by Sy1-5 strands superimposed on cylindrical representations of the β-barrel they partially surround. (d-f) Similar representations for a Type 2A octamer model for which Sy5,6,8 form the inner barrel and Sy1-4 form the outer barrel. (g,h) TEM images of HNE-αSOs and DHA-αSOs (Enlarged and modified from van Diggelen et al. [18]). We superimposed red and yellow circles with outer diameters of 6.0 nm and 7.1 nm predicted for the 2^nd^ β-barrel of α-Syn models of a hexamer and octamer around lighter shaded regions.

Unlike models described above, the concentric β-barrels models that follow have central pores. The ratio of strands/monomer of the inner barrel to the surrounding barrel decreases as the number of strands increases: for the tetramer described above the ratio was 2/5, for the hexamers described below it is 3/5, and for octamers to dodecamers it is 3/4. These pores may act as lipid and fatty-acid binding sites; they are sufficiently large to surround alkyl chains and their entrances contain positively charged lysines that should interact favorably with negatively charged head-groups. It is difficult to predict based on the data whether the smaller HNE-oligomers are more consistent with Type 2A (Fig. 8 a-c) hexamers or Type 2P hexamers described later in the annular protofibril section because the Nt-NAC barrel region of both models have the same dimensions. The Type 2A model has more antiparallel β-strands but the Type 2P models are more consistent with the absence of A11 binding. The hexamer model differs from our other models in that Sy7 forms a continuous β-strand that is part of a β-barrel and Sy7 and Sy8 form a β-hairpin. Perhaps the absence of Sy7 in β-Syn explains why its oligomerization is unaffected by fatty acids [58]. The data for the DHA oligomer are more consistent with a Type 2A octamer (Fig. 8 d-f).

#### Oxidized Dodecamers

In an oxidizing environment of Fe(II) and oxygen, N-terminally acetylated α-Syn forms oligomers with a substantial amount of right-hand twisted antiparallel β but no parallel β secondary structure [59]. EM images indicate that these oligomers aggregate with the smallest apparent components having a diameter of ∼ 9 nm. Fig.9 illustrates a Type 2A dodecamer tri-β-barrel model of α-Syn consistent with these dimensions. The Nt and NAC domains of each monomer resemble those of the octamers of Fig. 8. However, in the dodecamer model the Sy5 and the NAC domain forms the middle β-barrel. As in some of our other models, each Ct domain forms a 4-stranded antiparallel β-sheet; but, unlike other models, these sheets form two inner β-barrels. These 24-stranded antiparallel β-barrels each have 6-fold radial symmetry and stack end-on with 2-fold perpendicular symmetry. Most of the negatively charged side-chains of the Ct-barrels either extend inwardly into the central pore where they may bind to Fe^2+^ ions, or are located at the ends of the barrels that are exposed to the aqueous phase or can bind to lysines in the connections between strands of the central barrel. The only outwardly oriented negatively charged side-chain near the central regions of the barrels, E130, may salt bridge to the inwardly oriented H50 in Sy5 of the middle barrel. K102 and E104 near the beginning of the first Ct strand may bind to D98 and K96 near the end of the Sy8 strand. If S/N of the inner, middle, and outer barrels are 1.0, 2/3, and 0.5, the pitches of the pleats will be about the same for all three barrels, facilitating intermeshing pleats. The theoretical gap distances between the walls of adjacent barrels should be ∼ 0.75 and 0.85, consistent with the small sizes of the side-chains (primarily G, A, V, S, and T) that form the interacting pleats. This dodecamer model has six-fold radial symmetry with a relatively hydrophobic outer surface in the region of 2-fold perpendicular symmetry occupied by Sy4 strands (yellow-green). These features may facilitate aggregation into hexagonal arrays (Fig. 9c). The red circles in Fig. 9 with the diameter predicted for the dodecamer illustrate the consistency of this hypothesis with the EM images.

**Figure 9.**
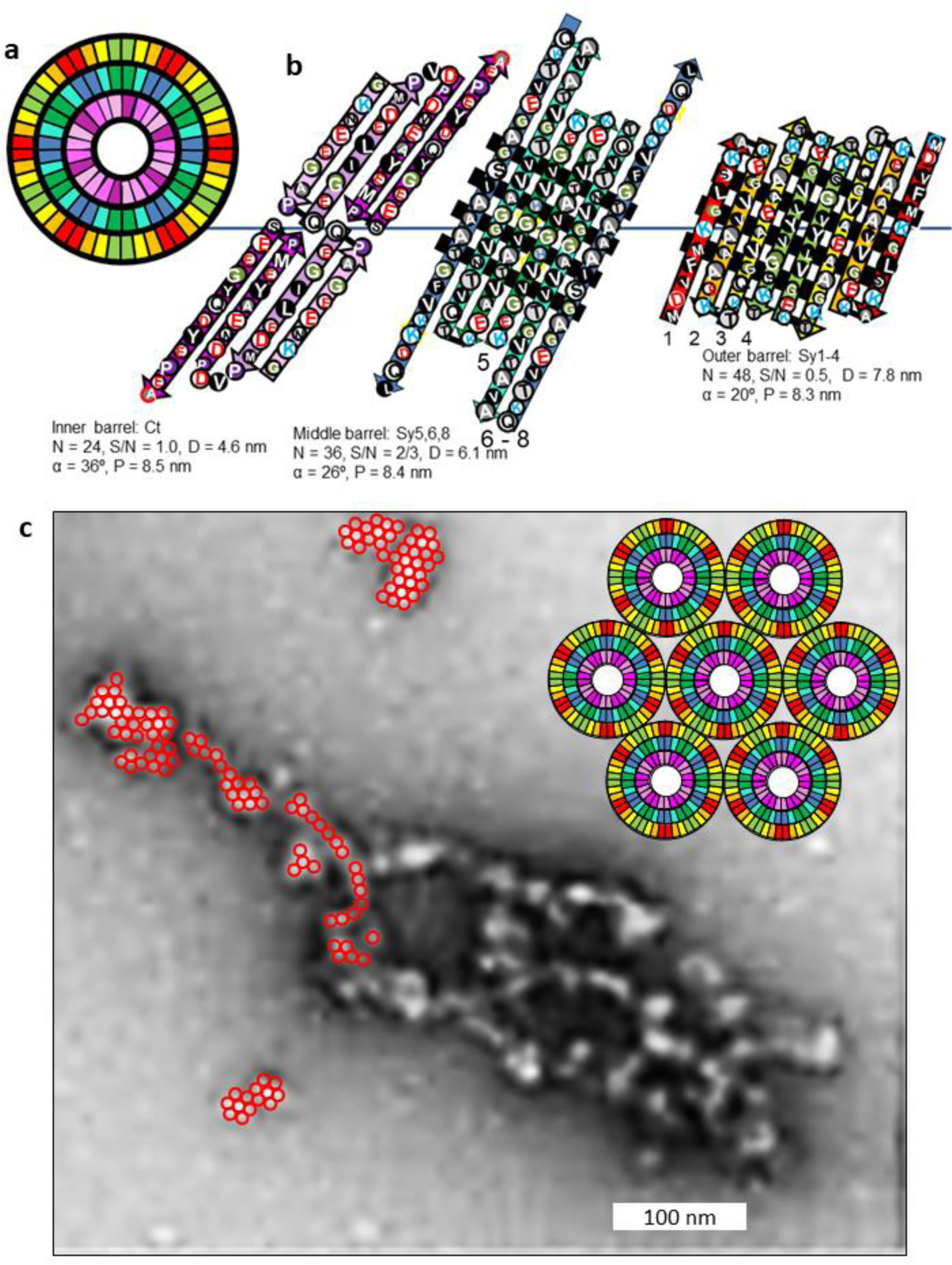
A Type 2A dodecamer model for α-Syn in the presence of Fe II and O_2_ and EM images of the assemblies. (a) Wedge representation of the model. (b) Flattened representations of two antiparallel subunits of the three concentric β-barrels. The inwardly oriented pleats of the middle and outer barrel are highlighted with black rectangles in the background. The Ct domains form two β-barrels, each with 6-fold radial symmetry, that stack end on with 2-fold perpendicular symmetry. (c) EM images [59] of antiparallel α-Syn oligomers. (Adapted with permission from Abeyawardhane DL, Fernandez RD, Murgas CJ, et al. Iron Redox Chemistry Promotes Antiparallel Oligomerization of alpha-Synuclein. J Am Chem Soc. 2018 Apr 18;140(15):5028-5032. Copyright 2018, American Chemical Society). The red circles superimposed on portions of the image have a diameter of ∼ 9 nm as predicted by the dodecamer models that are enlarged ten-fold in the upper left and arranged in a hexagonal array.

### Type 2P oligomers

#### Tetramers

With time α-Syn oligomers tend to transition from antiparallel to parallel β-structure [3]. The β-barrels of Type 2P models are parallel within the subunits and antiparallel between the subunits. The core of the 2P tetramer model is an 8-stranded β-barrel formed by Sy2 and Sy4 (Fig. 10). It is surrounded by a 16-stranded β-barrel formed by Sy1,3,5, & 7b-8. This barrel may be surrounded by a 3^rd^ 24-stranded antiparallel β-barrel formed by Sy6-7a and five Ct strands; however, if present this highly tentative outer barrel may be unstable and/or relatively disordered.

**Figure 10.**
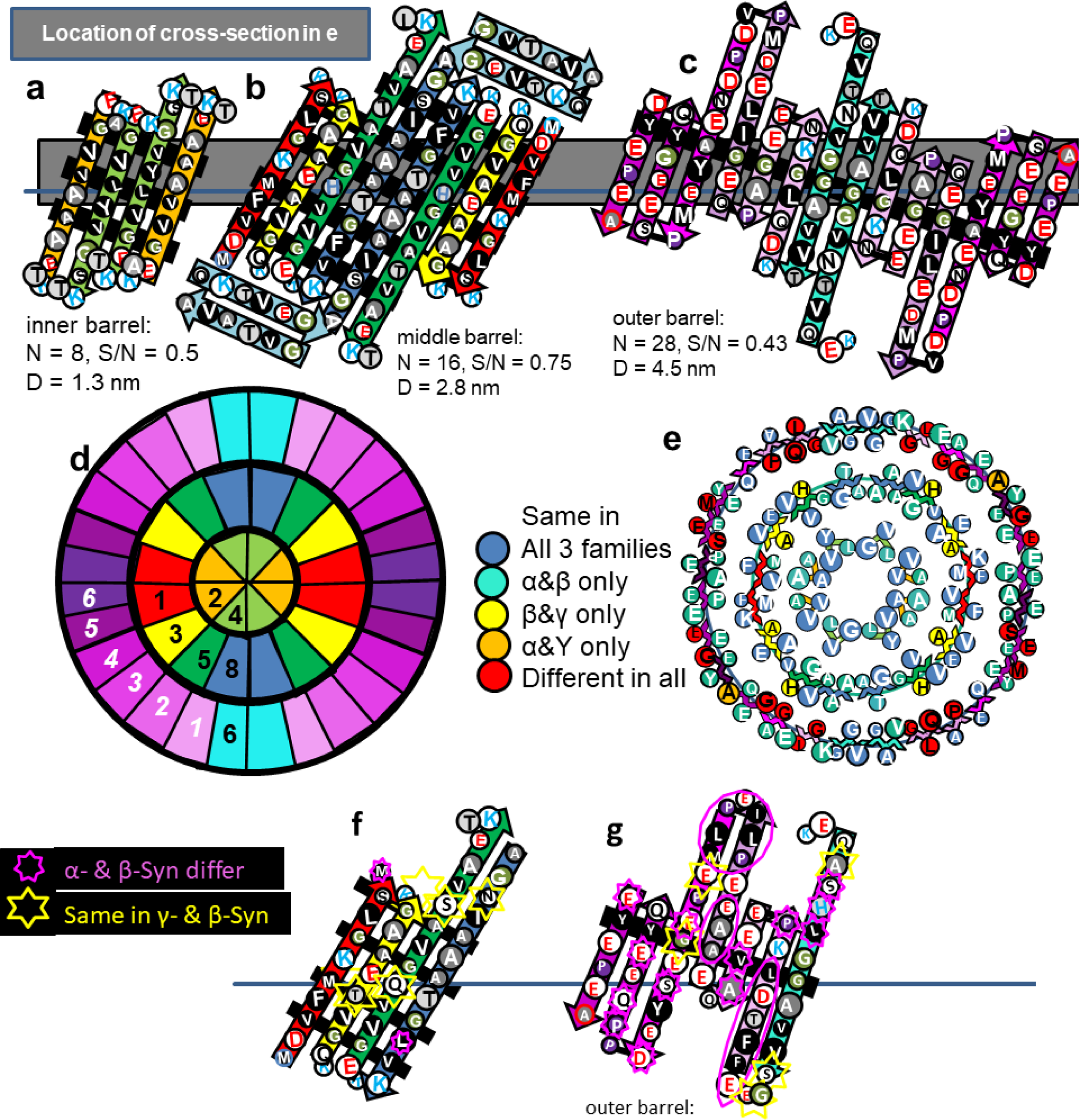
Type 2P model of an α-Syn tetramer with (a) Sy2 and Sy4 forming the core 8-stranded parallel β-barrel, (b) Sy1,3,5, & 8 forming the second 16-stranded parallel β-barrel, and (c) Sy7 + the Ct domain possibly forming an outer 24-stranded antiparallel β-barrel. Residues of the flattened representations are colored as in previous figures. (d) Wedge representation of a central cross-section showing proposed relative positions of β-strands. The Ct segments of the wedge representation are numbered in white and italicized. (e) A central cross-section illustrating degree of conservation of residues. The sequences of the inner two barrels are almost identical for the α-Syn and β-syn families and where they differ on the exterior of Sy3 and Sy5 of the middle barrel they are the same in the β-syn and Υ-Syn families. (f & g) Models of one subunit of the middle and outer β-barrels for β-Syn and positions where residues differ from α-Syn.

This model was developed to maximize interactions among and burial of apolar residues that are identical in α- and β-Syn sequences and to maximize the amount of parallel β-structure. (Reversing positions of Sy6 and Sy8 would simplify the 2P model; however, doing so would reduce clustering of the conserved apolar residues and the amount of parallel β-structure.) No residue differences occur in the putative β-strands of Sy2 and Sy4 or in all but one (A27T) inwardly-oriented side-chains of Sy1, Sy3, Sy5, or Sy8 β-strands that interact with those of Sy2 or Sy4, consistent with the hypothesis that this motif is conserved in both families. The second β-barrel of this model has four fewer β-strands and its diameter is ∼0.4 nm less for the 2A tetramer model of Fig. 7, making the side-chain packing more compact in the 2P model and the distance between the walls of the two β-barrels only 0.7 nm. This tight packing is feasible because small apolar side-chains dominate the area between the inner and outer β-barrels; each monomer has seven glycines, seven alanines, six valines, two serines, one methionine, and one tyrosine side-chain between the barrels. If Type 2P tetramers are essential for the formation of some functional α- and β-Syn assemblies, then tight and precise packing of the hydrophobic core may explain why even conservative substitutions do not occur between the two families for these interacting residues.

The only charged side-chains of the putative Nt β-strands face outwardly on the middle β-barrel: the negatively charged E20 of Sy3 is positioned between positively charged K6 of Sy1 and H50 of Sy5. This outwardly oriented pleat of charged side-chains is flanked on each side by a pleat of hydrophobic side-chains.

The tentative model of the outer barrel was chosen for several reasons: (1) The inwardly oriented central pleat of adjacent subunits has eight adjacent glycines flanked by an alanine and two tyrosines on each side. The small residues of the central region should reduce side-chain clashes between the barrels. (2) The pitches of the pleats are about the same for the middle (4.8 nm) and outer barrels (4.2 nm). (3) Most prolines are located in the second position of β-turns, a frequently-observed location for prolines. (4) Indels in the α-Syn and β-Syn sequences occur in turn regions between the β-strands where they should have minimal effect on the structures. (5) Four consecutive residues (GGAV) within the Sy6 strand interact with their counterparts on an axis of 2-fold symmetry and conserved apolar residues of two interacting S8 segments of the middle β-barrel. (6) Seven of the last eight residues of the Ct domain are identical in α-Syn and β-Syn sequences and self-associate at the other 2-fold axis and cover conserved portions of two interacting Sy1 strands. (7) The rest of Sy6 and most of the Ct domain are poorly conserved and interact with less conserved portions of the middle barrel (e.g., H50Q) or the aqueous solvent. (8) The putative outer layer strands are positioned to allow connections of Sy5 to Sy6, Sy6 to Sy7, and Sy8 to Ct1. (9) The Sy7 β-hairpins are on the top and bottom of the assembly where they are exposed to solvent and should have little effect on the rest of the structure.

The model of Fig.10 f and g for the β-Syn tetramer differs somewhat; e.g., the Sy7 segment is deleted and there are some insertions in the first portion of the Ct domain. These indels should not affect the rest of the structure dramatically because they do not occur within the putative β-barrels. The outer barrels of both models have the same number of strands and a central inwardly oriented pleat (black background) composed of apolar and relatively conserved residues sould fit between outwardly oriented pleats of the middle barrel.

Excluding the tentative outer barrel of the 2P model, all trimer and tetramer models described above have diameters consistent with the bead images in Fig. 4.

#### Octamers with solid cores

Two Type 2P tetramers may merge to form an octamer with a similar concentric β-barrel structure and a highly conserved apolar core (Fig. 11); the tetra and octamer models each have an 8-stranded inner β-barrel, a 16 stranded 2nd β-barrel, and a 24-stranded third β-barrel. The octamer model has an additional, but tentative, fourth β-barrel formed by Ct domains. Although the relative positions of Nt strands within and between the monomers are the same in both models, these segments comprise the 2^nd^ and 3^rd^ barrel in the octamer. A major difference is that the highly conserved hydrophobic Sy8 strands comprise the innermost 8-stranded β-barrel in the octamer. S/N must be 1.0 for the core barrel to have 4-fold radial symmetry; thus, strands of all but the outer barrels are tilted by 36° and the gap distances between barrels is ∼ 0.76 nm. Of all models presented here, these tetramer and octamer ones are most consistent with the patterns of conservation among the three synuclein families. The strands of the first three β-barrels all have at least six residues that, with the exception of a few outwardly oriented residues of the third barrel’s Sy1,3,5 segments, are apolar or have only hydroxyl side chain moieties. Almost all β-barrel residues interior to the wall of the 3^rd^ barrel are small and identical in α-Syn and β-syn.

**Figure 11.**
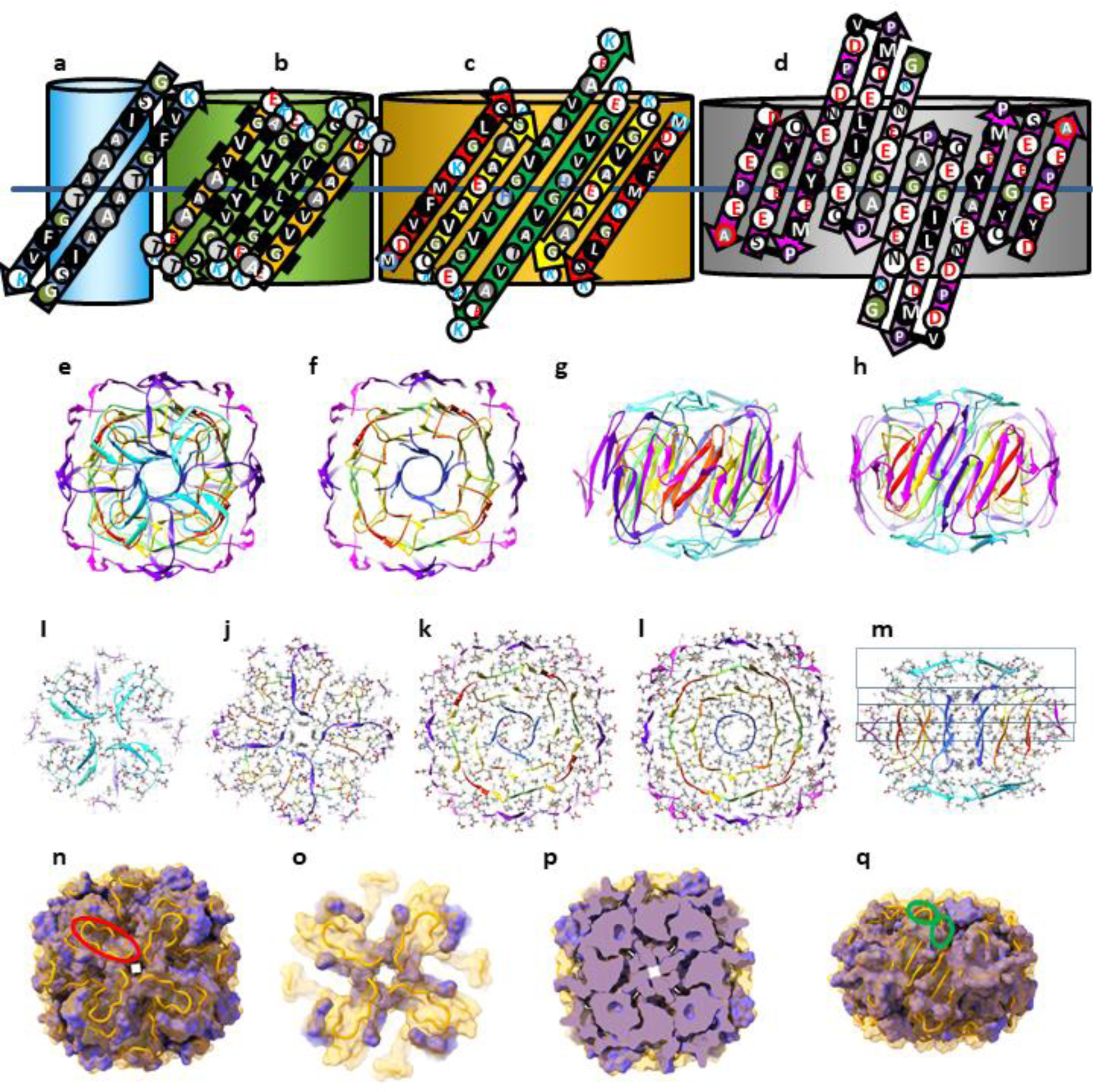
Models of an α-Syn Type 2P octamer with a small central pore. (a-d) Flattened representations for two α-Syn subunits of the four putative β-barrels superimposed on cylinder representations of each barrel. Sy6, Sy7, and Ct1 segments are not part of the core barrels and models of these segments are more speculative; they may be relatively disordered and are not integral components of the model. (e-h) Ribbon representation of an atomic-scale model colored by the rainbow beginning with red and ending with purple. (e) Top view down the axis of the pore. Sy6 and Sy7 segments that connect Sy5 to Sy8 are cyan and the Ct1 segment that connects Sy8 of the inner barrel to Ct2 of the outer barrel is blue-purple. (f) Clipped view of the four concentric β-barrels. (g) Side view down the 2-fold axis of symmetry between the red Sy1 strands. Portions of the outer barrel have been clipped away. (h) Side view rotated by 45⁰ relative to (g) viewed down the axis of 2-fold symmetry formed between the purple Ct6 and green Sy5 strands. (I-m) Cross-sections illustrating side-chain (colored by element) packing between the barrels: (i) Top view, (j) one layer down, (k) two layers down, (l) central layer, (m) side-view showing the positions of the layers. (n-q) Illustration of the surface of conserved core (purple) formed by residues that are identical in α-Syn and β-Syn. The orange tubes are the back-bone trace of residues that are not identical in the two sequences and for which the surfaces are not shown. (n) Top view; one of the Sy7 segments that is deleted in β-Syn is encircled in red. (o) Cross-section of the top segments connecting strands of the β-barrels. The conserved purple portions are at the ends of Sy5 and Sy6. The rest of the connectors are not conserved. (p) Cross-section of the mid-region illustrating dense packing of the conserved core (purple). (q) Side view with the same orientation as in (g).

Poorly conserved Sy6, Sy7, and the Ct1 segments extend above and below the core tri-β-barrels where they connect strands of different β-barrels.

We developed atomic scale models of the octamer in order to explore and illustrate the feasibility of the structure in more detail (Fig. 11 e-q). The backbone and secondary structures of the model are illustrated by rainbow-colored ribbons (e-h). Cross-sections showing side-chains colored by element demonstrate relatively dense side-chain packing (i-m). Residues that are identical in α-Syn and β-Syn cluster together to form a densly-packed core. Residues that differ in the two sequences and where indels occur are located primarily on the surface and/or interact with other non-conserved residues (n-q). Charged side chains of the Nt and NAC domains can all form salt bridges.

#### Annular and tubular protofibrils

Amyloid oligomers typically form protofibrils which in turn morph into fibrils. In some conditions annular protofibrils of α-Syn and mutant A30P and A53P α-Syn form first then combine to form tubular protofibrils [4]. We have developed Type 2P hexamer, octamer, decamer, and dodecamer models of annular protofibrils with diameters consistent with EM images of annular protofibrils (Fig. 12).

**Figure 12.**
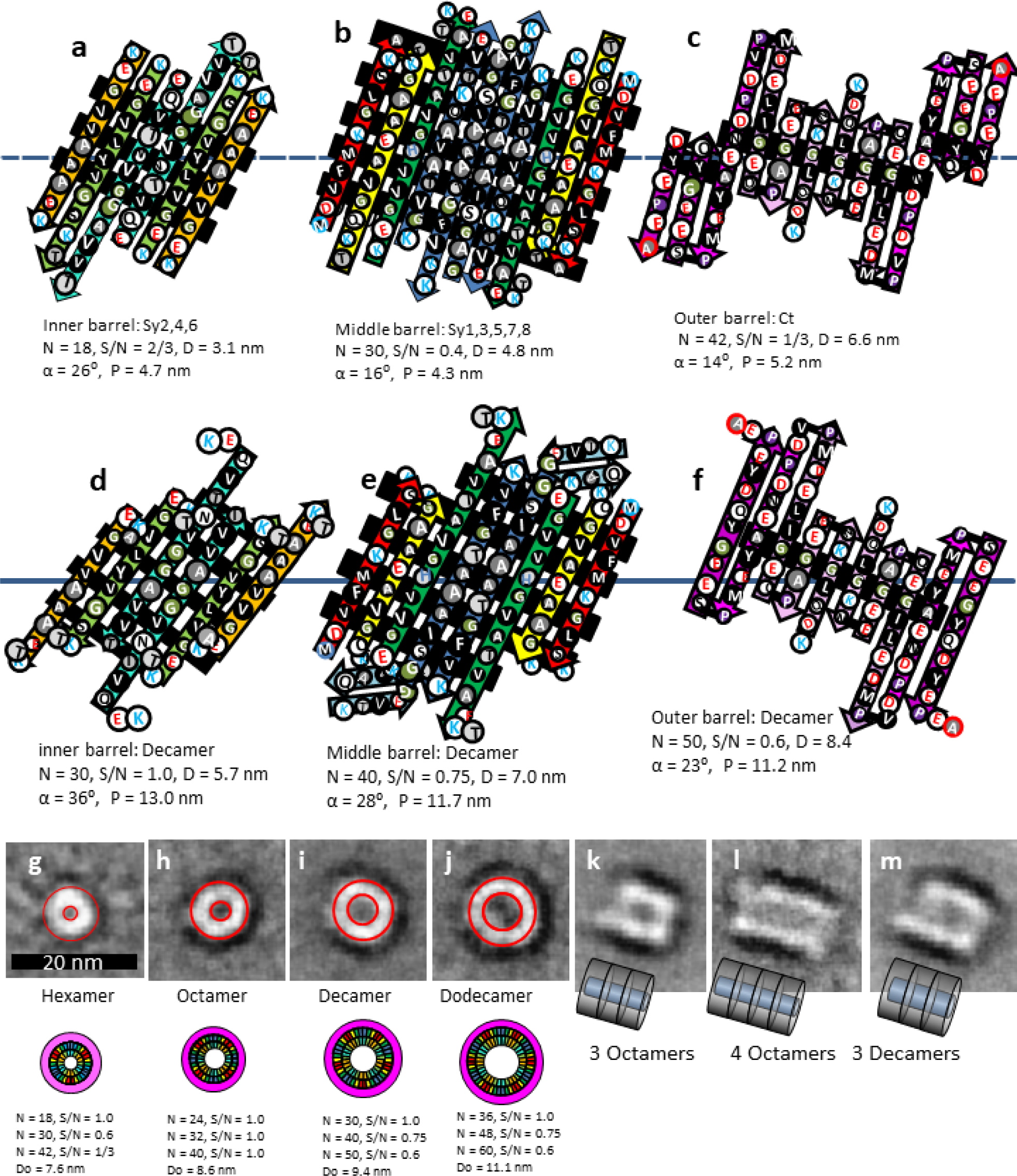
Models and images of mutant α-Syn annular and tubular protofibrils. (a-f) Flattened representation of two adjacent subunits of Type 2P models for a hexamer (a -c) and a decamer (d-f). Parameters of the barrels are listed below the figures. Inwardly oriented pleats composed primarily of small apolar side-chains are highlighted with black backgrounds. (h-m) EM images of four sizes of (g) A30P and (h-j) A53T annular protofibrils, and wedge representations of our models of a hexamer, octamer, decamer, and dodecamer drawn to scale. The red circles on the images represent the outer and inner APF boundaries predicted by the models. (k-m) Side views of three A53T tubular protofibrils that develop from the annular protofibrils. Schematics represent our models of how the annular protofibrils may pack end-to-end to form the tubular protofibrils. EM images were copied with modification and permission from Lashuel et al. [4].

The general folding pattern for the A30P mutant hexamer (Fig 12 a-c) resembles the Type 2P tetramer of Fig.10 except that Sy7 forms a continuous β-strand that is a component of the middle barrel. There should be few side-chain clashes between the barrels. The gap distances between the barrels are 0.85 and 0.9 nm and the pitches of the pleats are similar for the three β-barrels so the pleats should intermesh well. One central inward pleat of the Ct domain has six consecutive glycines that should fit between pleats on Sy7 and Sy8 of the middle barrel that are composed of alanines. The central inward pleat of the middle barrel is composed of glycines, alanines and valines and should fit between pleats of the inner barrel.

The folding patterns for the A53T mutant octamers, decamers, and octamers are similar; Sy6 and Sy8 each contribute an additional β-strand/monomer to the inner and middle barrels; i.e., Sy2,4,6 comprise the inner β-barrel, Sy1,3,5,8 comprise a middle β-barrel, and the Ct domains that comprise an outer β-barrel of these models do not include Sy6 (Fig. 12 e-g). The number of strands per monomer increases by one for each concentric barrel. Sy7 is proposed to form a β-hairpin, but its location in the illustration is somewhat arbitrary; constrained primarily by the assumption that the folding pattern should be viable for β-Syn that has no Sy7 segment. The gap distances between barrels ranges from 0.6 to 0.9 nm for the four models. All three barrels of the octamer would have S/N values of 1.0 with gap distances of ∼0.76 nm between barrels. Tight packing between barrels can be maintained by reducing S/N values of the 2^nd^ and 3^rd^ barrels to 0.75 and 0.60 for the decamer (Fig. 12 a-c) and dodecamer models; yielding gap distances ranging from 0.65 to 0.84 nm between barrels. Tight packing between barrels of the decamer and dodecamer may be facilitated by intermeshing pleats because the pitches of the three barrels are almost the same and most side-chains of the interacting pleats are small (glycine and alanine for the central region of the central pleat). The proposed structure of the outer Ct barrel resembles those in Figs. 10 and 11.

The 2-fold perpendicular symmetry of Type 2 models is consistent with the end-to-end interactions proposed for how annular protofibrils form tubular protofibrils. Interactions among the putative Sy7 β-hairpins may facilitate interactions among the annular protofibrils to form the tubular protofibrils; β-Syn that lacks Sy7 does not form fibrils and the Sy7 segment appears to be essential to fibril formation [60].

#### High density lipoproteins. α-Syn and β-Syn

All three familes of Synucleins interact with sphingmylein to form high density lipoproteins [19]. These complexes are not toxic and will form with both native and mutants of α-Syn associated with early onset Parkinson’s disease. α-Syn and β-Syn lipoproteins assemble into stacks of disc-shaped complexes (Figs. 13 and 14). Both discs and individual oligomers are visiable in the EM images. The circular oligomers frequently cluster; the diameters of most oligomers in these clusters are consistent with our models of Type 2P tetramers and octamers if the Ct barrels are not included.

**Figure 13.**
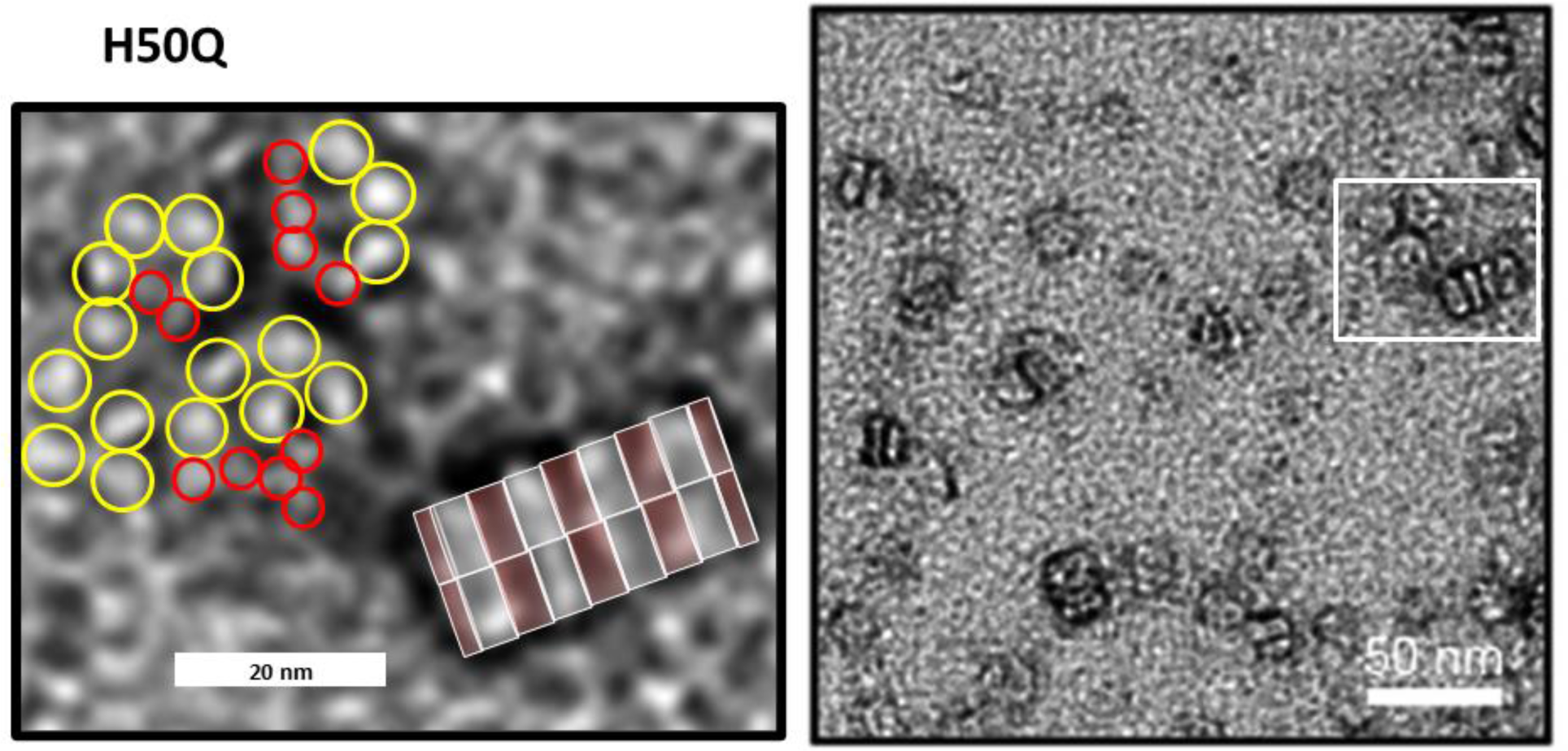
EM images of lipoproteins formed by α-Syn mutant H50Q. The region enclosed in the rectangle on the right is enlarged on the left. The red and yellow circles have diameters of 3.8 and 5.6 nm predicted by the tetramer and octamer models of Figs. 10 and 11 if the outer Ct barrel is excluded. The gray and pink transparent rectangles over the stack represent side views of octamer Nt-NAC and Ct domains, each pair is 7.5 nm apart. (EM images were modified from Eichmann et al. [19]).

**Figure 14.**
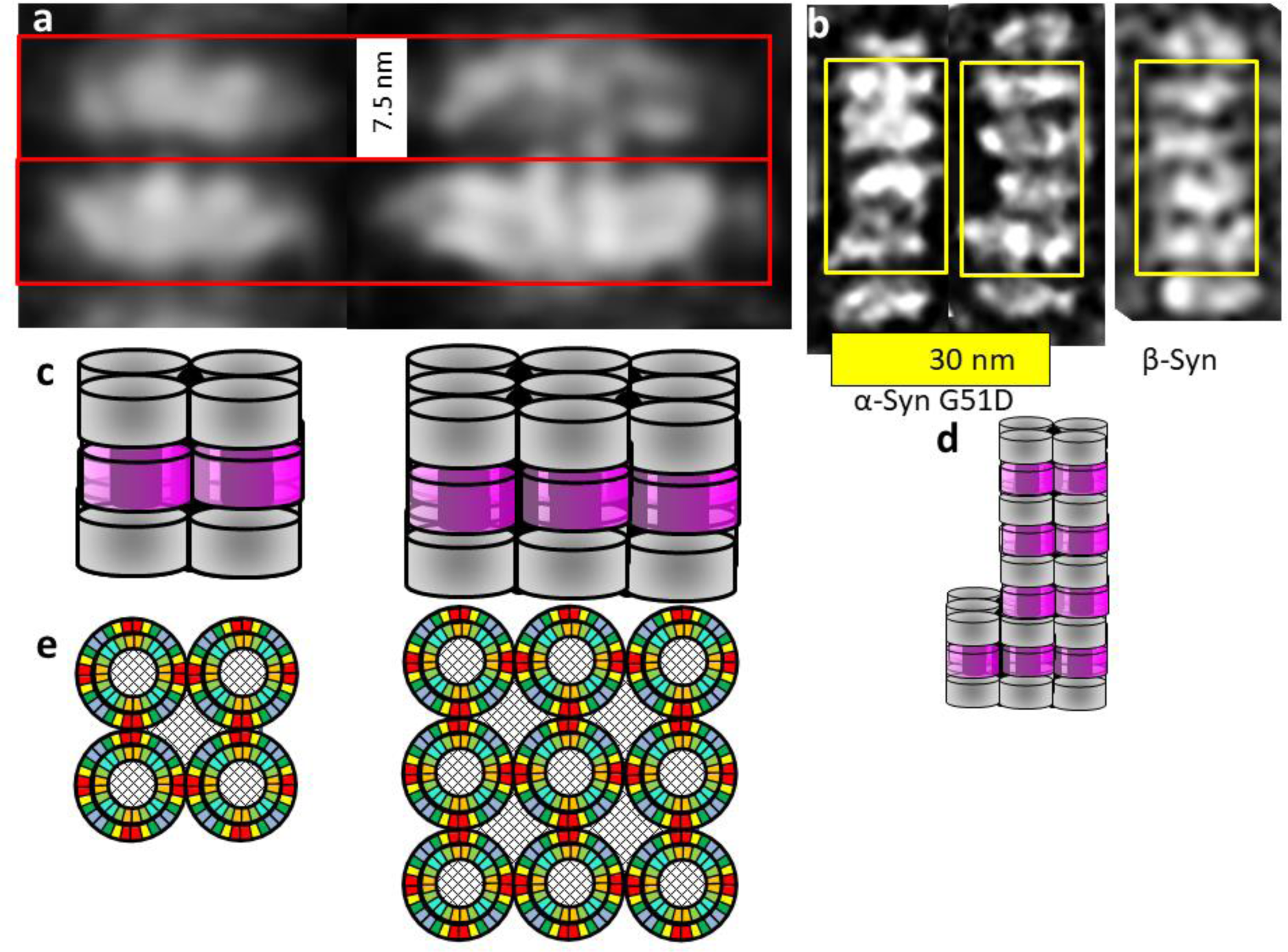
Images and models of α-Syn and β-Syn stacked lipoprotein discs. (a) Two sizes of α-Syn stacked pairs obtained by image averaging. (b) Individual images of relatively long stacks for both α- and β-Syn. (Images modified from original EM images of Eichmann et al. [19]). (c) Side views of schematics of our models. The darkly shaded regions in the middle of and between the cylinders represent lipid-filled regions. The gray cylinders represent concentric β-barrels formed by the Nt and NAC domains from two discs; the magenta cylinders represent 32-stranded β-barrels formed by interlocking Ct domains from adjacent stacks. (d) A schematic model for the central stack illustrating variations of the numbers of octamers in adjacent discs of the stack above. (e) Wedge cross-sections representations of the Nt-NAC barrel regions of the octamers within a disc. The checkered regions inside and between the octamers represent lipid alkyl chains.

The distance between discs is a relatively constant 7.5 nm, but diameters of the discs vary. Each appears to be comprised of smaller components, suggesting that the number of subassemblies per disc vary (Fig 14a & b). Disc pairs that have approximately the same diameter were grouped; image averaging with Adobe Photoshop was applied to each group to obtain more detailed images (Fig. 14a). These averaged images reveal vertical columns within and sometimes between the discs that are ∼ 3.5 nm apart.

We considered several models for the basic components of the α- and β-Syn discs. The Type 2P octamer model illustrated in Figs. 15 and Supplement S1 were the most consistent with dimensions obtained from the EM images. Sy2,4,6 form an inner 24-stranded β-barrel and Sy1,3,5,8 form a 32-stranded outer β-barrel. Four-stranded Ct β-sheets from adjacent octamers interlock to form a 32-stranded antiparallel β-barrel that connects octamers. The pore through the inner β-barrel is lined with apolar side-chains and is ∼ 3 nm in diameter. This region is proposed to surround sphingolipid alkyl chains. The entrances to this pore are surrounded by positively charged lysines of the signature connectors. These may interact with negatively charged phosphate moieties of the lipid head-groups. The negatively-charged Ct β-barrel is proposed to surround the positively charged portions of the sphingolipid head-groups. The calculated distance between discs based on these model is ∼ 7.6 nm, about the same as observed in the micrographs.

**Figure 15.**
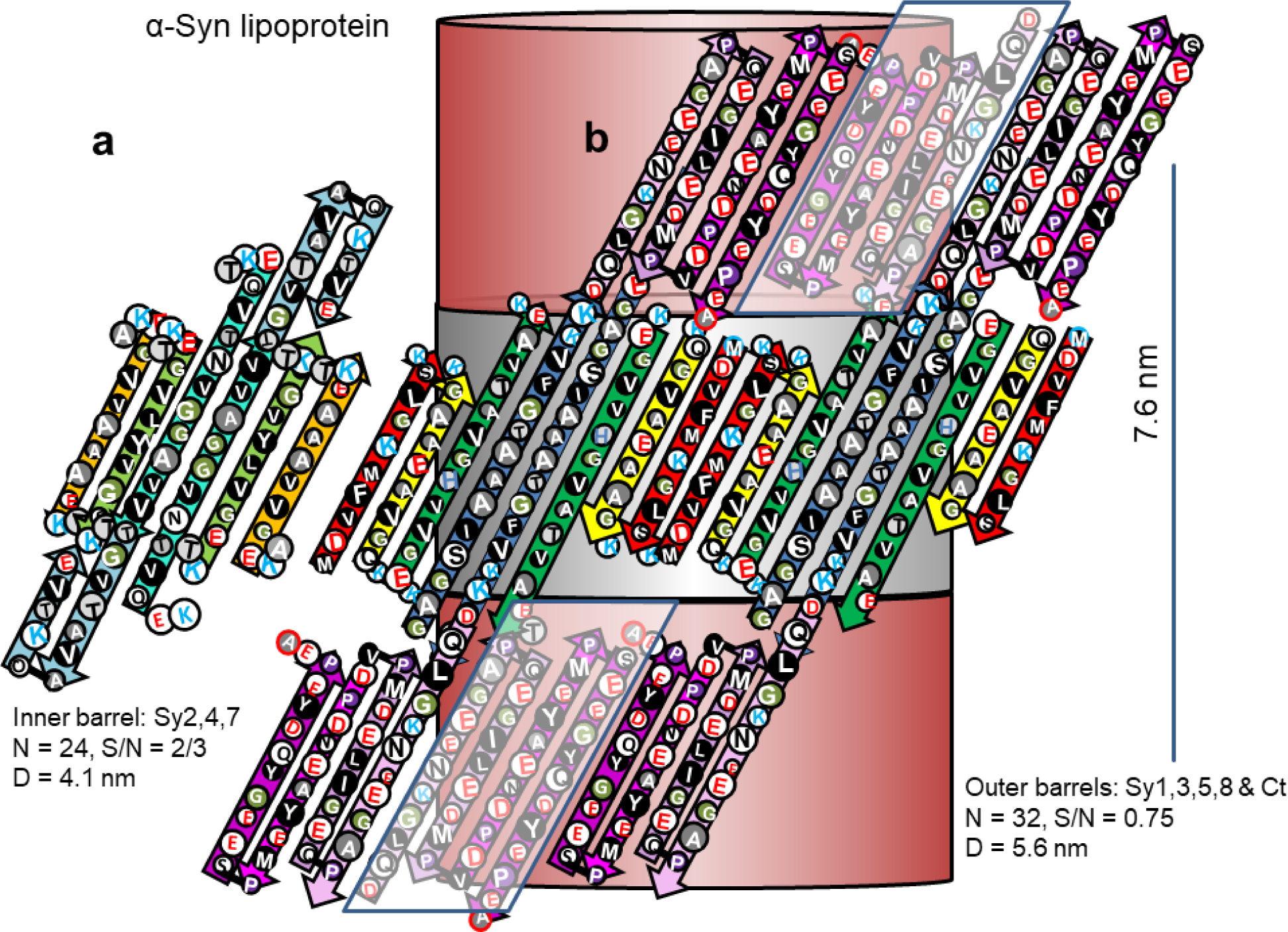
Flattened representations of the proposed α-Syn lipoprotein Type 2P octamer of discs. (a) Two subunits of the inner 24-stranded β-barrel formed by Sy2,4,6. (b) Four subunits of the 32-stranded outer β-barrels formed by Sy1,3,5,8. The Ct domains form 4-stranded antiparallel β sheets that extend in series with the outer β-barrels. The schematic for the Ct barrels on each end of the octamers includes Ct domains from adjacent octamers, enclosed in parallelagrams and in lighter shades. The cylinders in the background represent the β barrels that the strands wrap around. The calculated distance between adjacent discs is ∼7.6 nm.

The next challenge was to account for the lightly-shaded vertical columns that are ∼3.5 nm apart in the image-averaged discs (Fig. 14a). The Type 2P octamer structure has four-fold radial symmetry and 2-fold perpendicular symmetry. Adjacent octamers within the same disc likely form a square lattice in which all contacts are identical and occur at regions of 2-fold symmetry (Fig. 14 e). In an aqueous environment these contacts would likely occur in the most hydrophobic regions; i.e., between antiparallel pairs of Sy8 strands. However, in a hydrophobic environment the contacts would likely occur between more polar segments; i.e., at antiparallel pairs of Sy1 strands and adjacent Sy3 strands. The latter case would occur if lipids occupy the regions between the octamers, as illustrated in Fig. 14e. Positively charged lysines at the ends of the Sy5-8 strands could interact with negatively charged components of inter-octamer lipid head groups, and negatively charged side-chains on the exterior of the Ct β-barrels could interact with positively charged components of these lipids. This arrangement creates two types of lipid columns (intra-octamer and inter-octamer). If the diameter of the octamers is ∼7 nm, these columns would be ∼ 3.5 nm apart.

In spite of the absence of the Sy7 segment and substantial sequences differences in Ct1, Ct2, and Ct3, β-Syn discs can be modeled with the same general folding motif as α-Syn (Supplement Fig. S1). The requirement that both families have the same basic structure was used to constrain the end of Sy6 to be proximal to the beginning of Sy8 and that the Sy7 β-hairpin not be an integral component of the α-Syn β-barrels. Ct1 and Ct2 of β-Syn have insertions in the same general spatial region as occupied by Sy7 in the model of α-Syn.

In summary, the structures predicted by these models are consistent with the dimensions of the stacked discs observed in micrographs, and are well designed to bind lipids with zwitterionic head-groups. These non-toxic structures may represent a major functional role for these synucleins in regulating and transporting lipids.

#### High density lipoproteins: γ-Syn

Sphingomylein-γ-Syn lipoproteins form elongateded assemblies (called filaments here) rather than stacked discs [19]. Most of these filaments are isolated; however some occur as parallel pairs. These pairs helped us propose that two sizes of filaments occur with diameters of ∼ 7 - 8 and 12 nm (Fig. 16). Unfortunately, it is more difficult to assertain the dimensions and periodicity of these filaments than for the stacked-discs.

**Figure 16.**
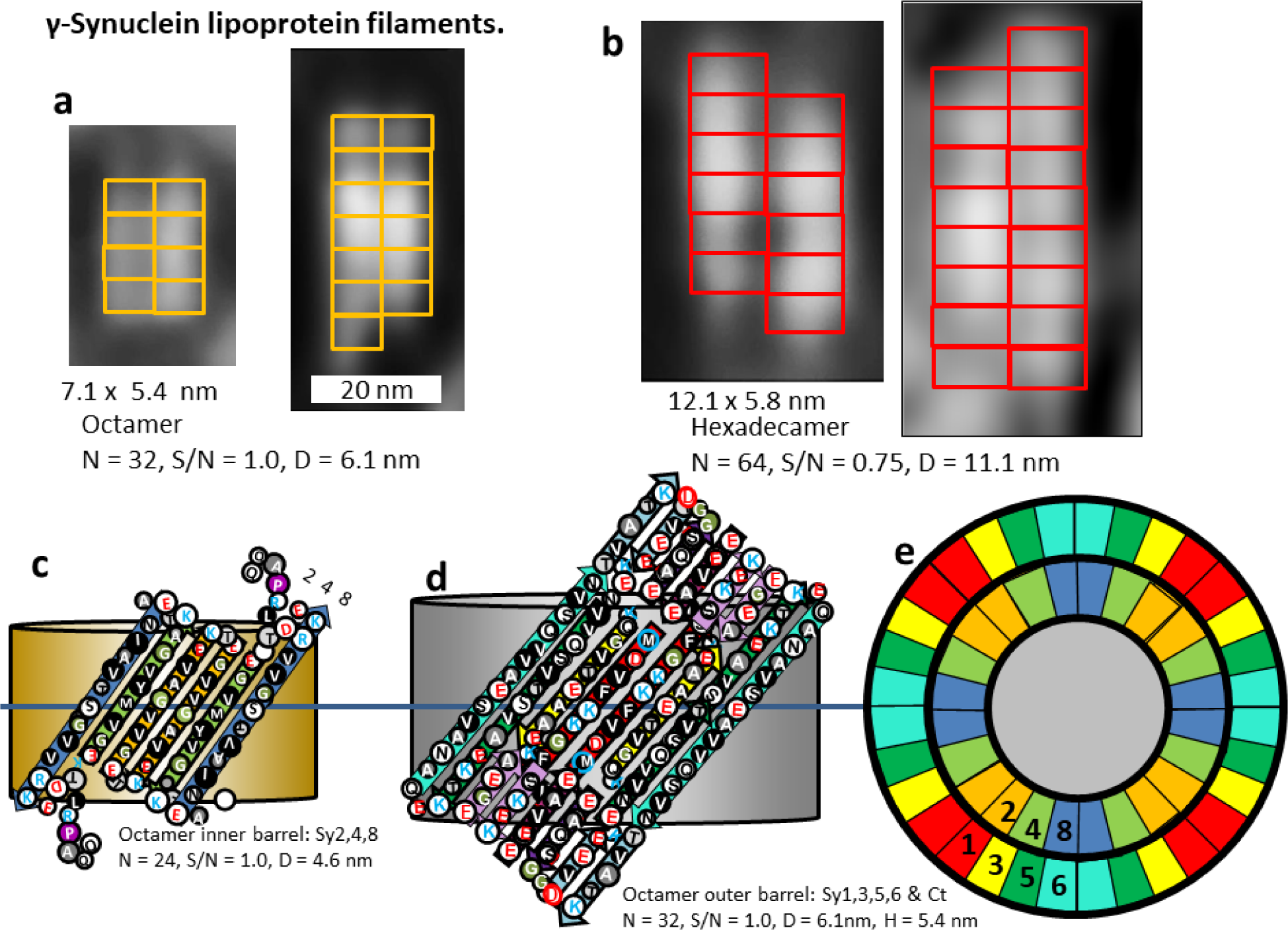
Negatively stained EM images of γ-Syn filament pairs and schematics of our models. (a) Smaller filaments. Image on the far left is averaged from 51 pairs and that on the right is an individual filament pair. The yellow rectangles represent models of Υ-Syn octamers stacked end-on. (b) Averaged and an individual image for larger filaments. Red rectangles represent models of hexadecamers stacked end-on. (c). Flattened represention for two subunits of the octamer inner barrel formed by Sy2,3,8. (d) Flattened representation for two subunits of outer 32-stranded β-barrels formed by Sy1,3,5,6 and by the Sy7 β-hairpin connecting Sy6 to Sy8, and four short β-strands of the Ct domain. The last two Ct strands could be from the same octamers as illustrated or from adjacent octamers. These barrels would stack end-on to form the outer walls of the filaments. (e) Wedge representation of a central cross-section of the two concentric β-barrels. The central region would be filled with lipid alkyl chains. The β-barrel parameters of a hexadecamer would be N = 48, S/N = 1.0, D = 9.2 nm, P = 21 nm and N = 64, S/N = 0.75, D = 11.1 nm, P = 19 nm.

The negatively charged Ct domain of γ-Syn is substantially shorter (most of Ct1 and Ct2 are missing) than those of α- and β-Syn and the sequences are difficult to align unambiguously. In this model four short segments of the Ct-domains interact with Sy7 β-hairpins and portions of Sy5 and Sy6 to form a negatively charged region of the pore that interacts with positively charged components of sphingomylein headgroups. The location of the last two Ct strands could depend upon interactions with other octamers; if there is no interaction they could have the positions indicated in the figure; if there is end-to-end interaction they could interact with Sy7 hairpins of the adjacent octamer. This region of the structure has some positively charged residues, but their side-chains are all oriented outwardly. As in the other models this Ct barrel is in series with Nt-NAC concentric β-barrels proposed to surround the lipid alkyl chains and acidic headgroup moieties. This model is consistent with sphingolipids binding inside the β-barrels but not between the β-barrels.

The γ-Syn lipoproteins also produce some substantially larger filaments (Fig. 16b). The diameter of the larger filaments is consistent with that predicted for assemblies with twice as many monomers as in the smaller filaments; *i.e.*, with a hexadecamer. S/N = 1.0 for the inner and Ct barrels in both models, however, the strands of the outer barrel are more tilted in the octamer than in the hexadecamer, making it more consistent with pleats-into-pleats packing.

## Equilibrium among Type 2 oligomers, protofibrils, and fibrils

Numerous studies indicate that α-Syn oligomers and fibrils exist in a dynamic equilibrium. An overview of our working hypotheses for this equilibrium among Type 2 oligomers and fibrils is given in Fig. 17. The oligomers of Fig. 4 with diameters consistent with our tetramer models tend to aggregate in strings. Antiparallel pairs of Sy5 β-strands comprise the longest and most hydrophobic regions on the exterior Type 2A tetramers; thus, they form the most probable regions of interactions between Type 2A tetramers (green strands in top left of Fig. 17). Portions of two Sy5 strands related by 2-fold symmetry pack tightly in some fibril structures [7,8,10] and most mutations leading to early-onset PD occur in Sy5. Thus it is plausible that interactions between Sy5 segments are vital to formation of functional α-Syn assemblies and that disruptions of these interactions favor formation of toxic assemblies. In contrast, antiparallel pairs of Sy8 strands of the NAC domains form the longest and most hydrophobic regions on the exterior of Type 2P tetramers (blue strands in Fig 17). Interactions of these segments occur at regions of 2-fold symmetry in Twister type fibrils [8]. Type 2P tetramers could be favored when concentrations are sufficiently high for most tetramers to form strings or clusters that bury the NAC segments. Additional monomers may bind to tetramers to form hexamers. Likewise, tetramers and hexamers may merge to form larger assemblies such as octamers proposed for lipoproteins and octamers, decamers, and dodecamers proposed for annular protofibrils and Fe II-O_2_ induced oligomers. Annular protofibrils may then interact to form tubular protofibrils, which slowly morph into fibrils. But interactions are not unidirectional; under some conditions fibrils can decompose to form oligomers that become smaller with time [2].

### Type 1 α-Syn Models: A Possible Toxic Pathway

Identification of which synuclein assemblies are essential and which are toxic may be vital for development of treatments. It is well established that an α-helical form of α-Syn binds to membrane surfaces and plays a vital role in the fusion of synaptic vesicles with the plasma membrane. However, interactions of α-Syn oligomers with membranes are not that simple: α-Syn adopts a multitude of membrane-bound conformations in unroofed cells and the Ct domain participates in membrane localization [61]. Using a microfluid setup that recapitulated the composition of the plasma membrane, Heo and Pinchet [23] found that α-Syn induced pores of discrete sizes and immobilized lipids on both the cis and trans sides of the membrane, indicating deep penetration inside the membrane. Quist et al. [21] used both electrical recording of membrane currents and atomic force microscopy to show that numerous amyloids, including α-Syn and its NAC domain, can form ion channels of multiple sizes in membranes. Kim et al. [22] also found that α-Syn oligomers formed well-defined conductance states in a variety of membranes and that the beta-structure of these membrane-bound oligomers differed from that of α-Syn fibrils. Parres-Gold et al. [24] used scanning ion conductance microscopy (SICM) to show that α-Syn oligomers induce morphological changes and formation of large transient pores in live SH-SY5Y neuroblastoma cells. Toxic α-Syn accumulation in SH-SY5Y is alleviated by membrane depolarization [62], suggesting that membrane interactions of α-Syn are related to toxicity in these cells. These interactions are affected by the lipid composition of the membranes. Binding of ganglioside and cholesterol appear to be involved in penetration of the membrane by α-Syn [26]. Based on results of nuclear magnet resonance spectroscopy Man et al. [63] concluded that cholesterol reduces the local affinity of the NAC domain to synaptic vesicles, supporting the role of disorder-to-order equilibrium of the NAC region. However, cholesterol-containing lipid nanodiscs accelerates oligomerization of α-Syn [64]. When the nanodiscs are made of DOPC, interaction of the NAC domain with the bilayer is increased; but when DOPC, DOPE, and DOPG all comprise the nanodiscs, binding of the NAC domain is not affected while binding of the Nt and Ct domains are inhibited.

Penetration of the alkyl phase of the membrane appears to depend upon the conformation of the α-Syn oligomer. Fusco et al. [65] analyzed two types of α-Syn oligomers of similar size that they called A and B. Type A interacts only with the surfaces of membranes, has little rigid secondary structure, and is not toxic. In contrast, segment 70-88 (red background in Figs. 18d and 19b) of the toxic Type B has a rigid β-secondary that penetrates the alkyl phase of the membrane. Most of this region is formed by Sy7 which is deleted in non-toxic β-Syn. Our working hypothesis is that their Type B oligomer is a Type 1 trimer in which the NAC domain forms a core six-stranded antiparallel β-barrel with the Sy7 segment in the turn region of the β-hairpin exposed.

**Figure 17.**
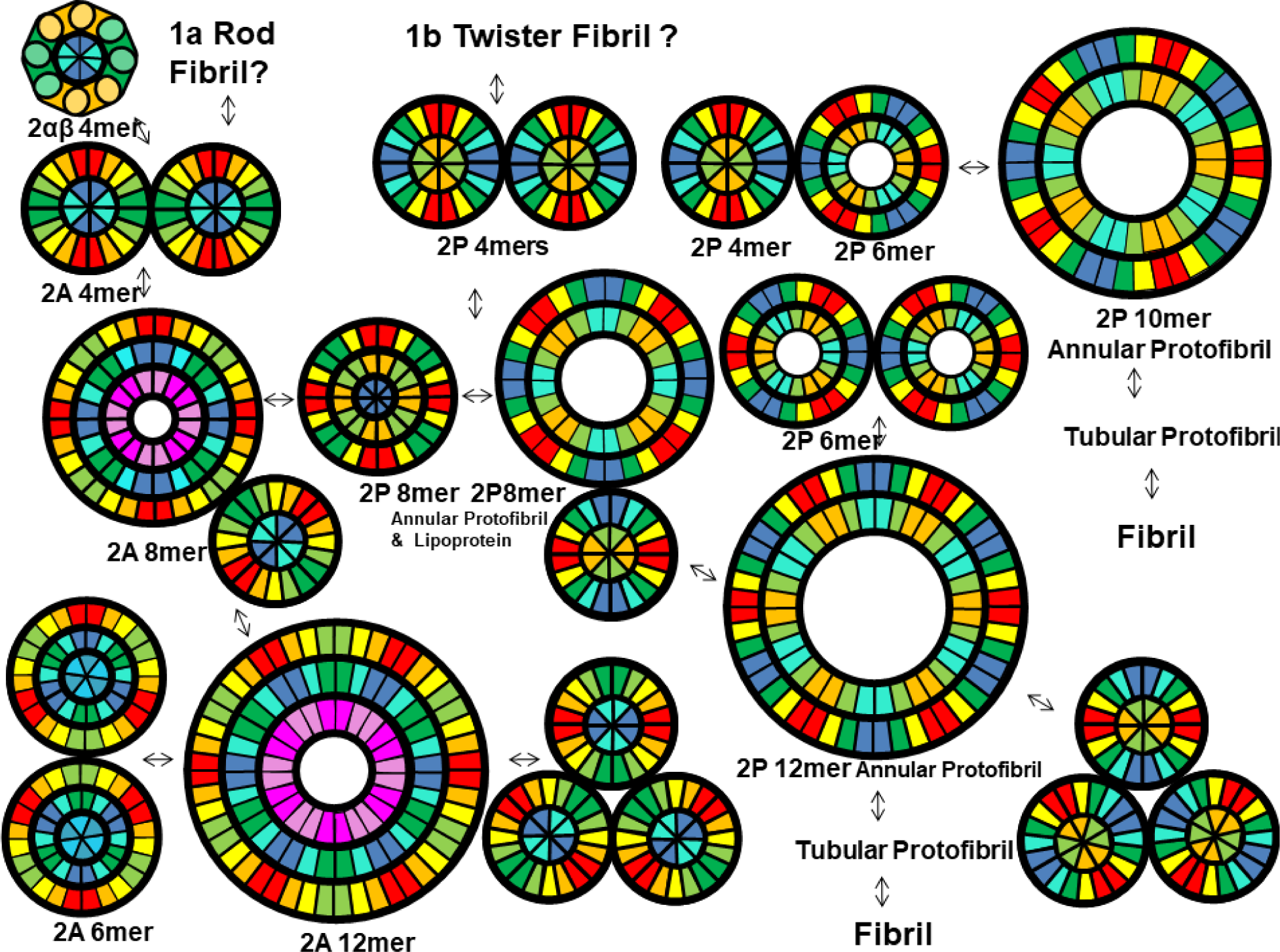
Hypothesis for dynamic equilibrium among Type 2 Synuclein oligomers, protofibrils, and fibrils. Wedge cross-section schematics were copied from previous figures.

**Figure 18.**
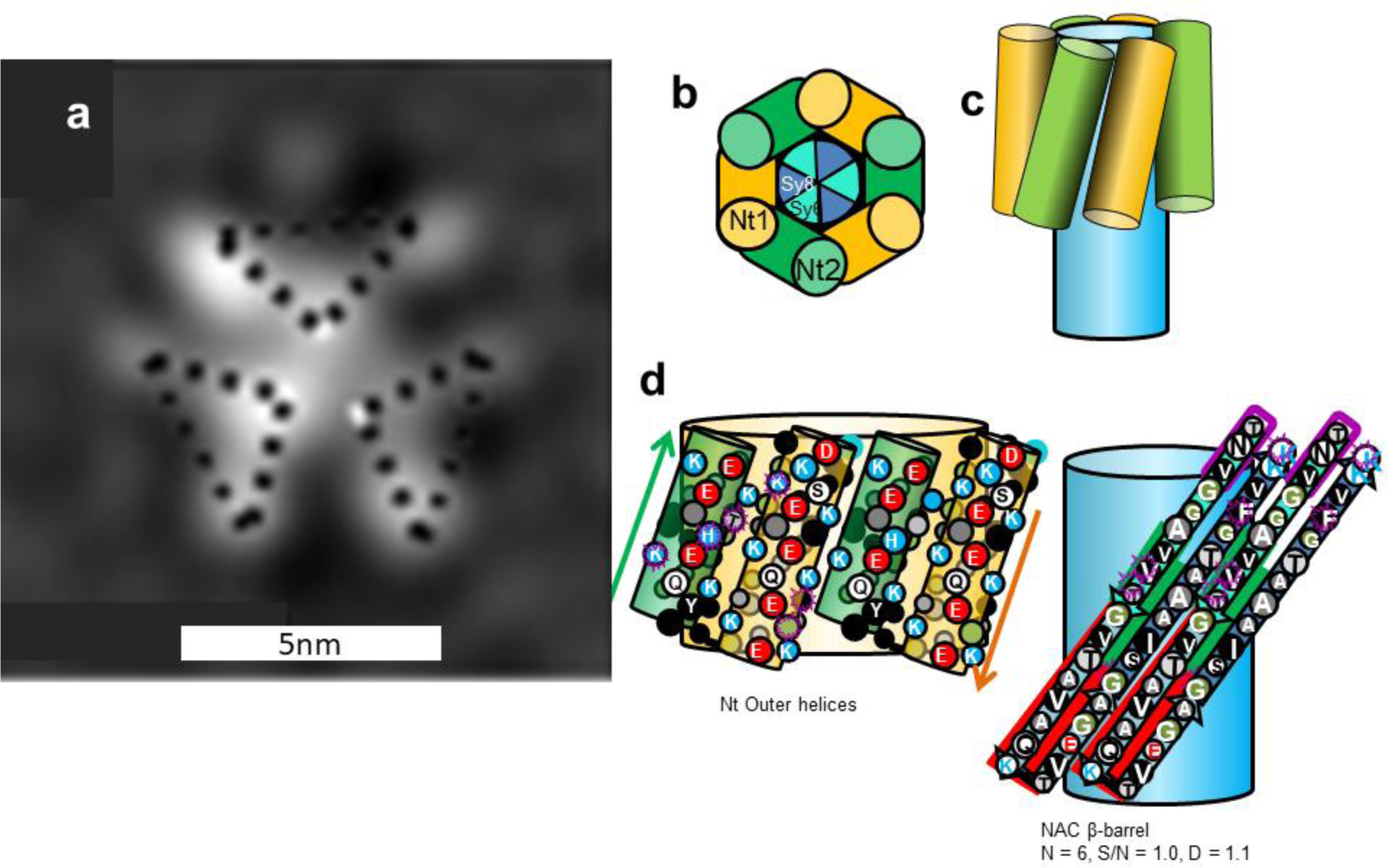
(a) EM image of a predominately α-helical trimer (Copied from Wang et al. [41]) and (b-e) a Type 1 α-β barrel trimer model. This model resembles the tetramer of Fig. 5 except that there is one less subunit and all subunits are oriented in the same direction. (b) wedge representation, (c) side view schematic of helices, (d) side view of two subunits of the outer Nt helices, (e) flattened representation of two subunits of a core 6-stranded antiparallel β-barrel formed by three NAC β-hairpins.

**Figure 19.**
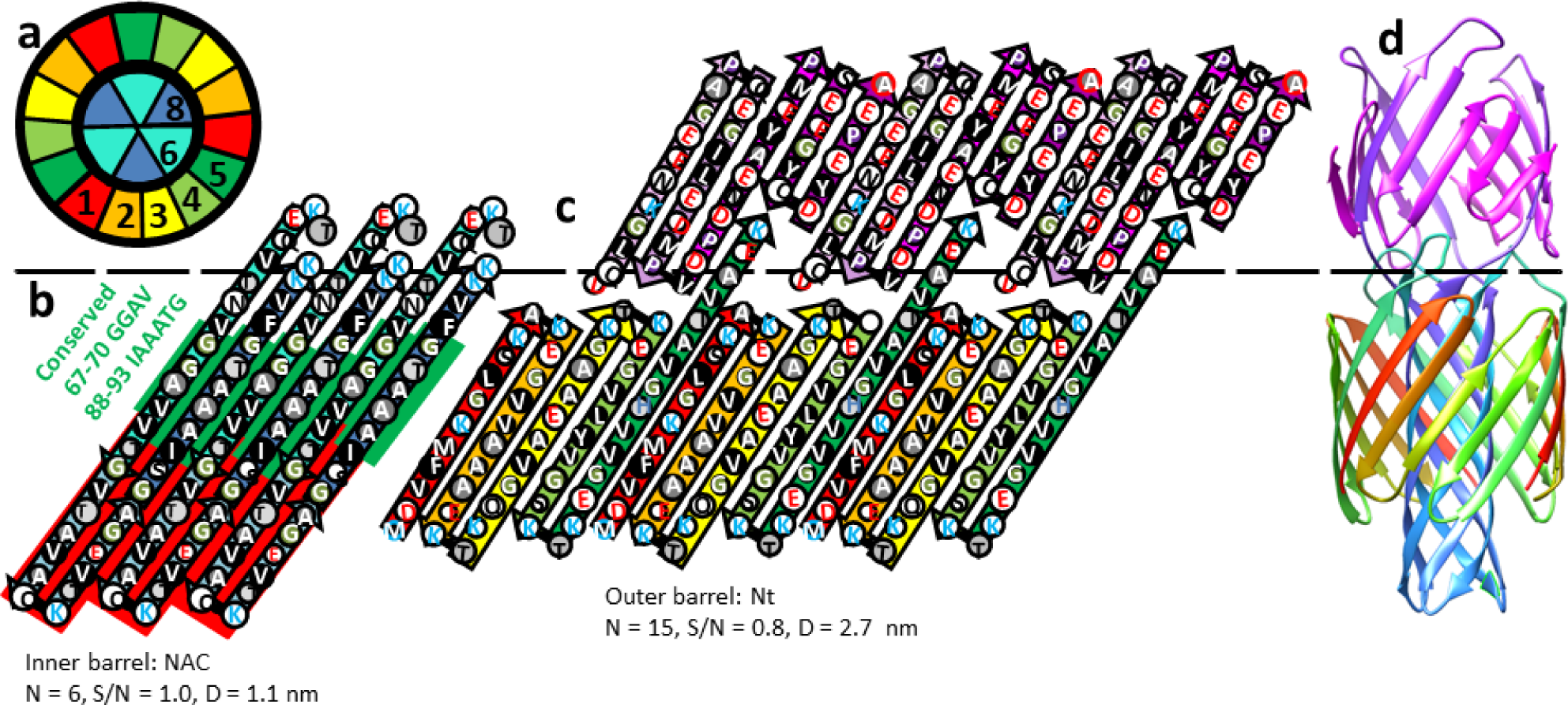
Concentric β-barrel models for Type 1A trimer. (a) Wedge schematic of a cross-section of the barrels. (b & c) Flattened representations of the inner and outer barrels of the trimer. The dashed line indicates the location where Sy5 and Ct1 of the outer barrels link to Sy6 and Sy8 of the inner barrel. Green and red backgrounds as in Fig. 5. (d) Ribbon representation of an atomic-scale model as viewed from the side. Ribbons are colored by the rainbow spectrum beginning with red at the N-termini and ending with magenta at the C-termini.

### αβ-barrel trimers

Type 2 structures must have an even number of subunits; however, Type 1 models can have any number. We have developed two types of α-Syn trimer models. Although α-helical tetramers of α-Syn have gained more attention, helical trimers have also been observed (Fig 18a). Helical trimers may transition to α-β-barrel trimers in which the NAC domains form a core six-stranded antiparallel β-barrel (Fig. 18b-e).

### Type 1A trimer

The α-β barrel trimer may transition to a concentric β-barrel structure in which the Nt domains form an outer 15-stranded antiparallel β-barrel that surrounds the core NAC β-barrel (Fig. 19). We examined this in greater detail by developing an atomic-scale model. The small size and apolar nature of the side-chains inside the core barrel and between the two barrels allows the gap distance between the two barrels to be small (∼ 0.8 nm) without disrupting backbone H-bonding within each barrel. The putative15-stranded Ct β-barrel structure is highly tentative; this domain is likely to be relatively disordered.

We suggest that if and/or when either of these structures interact with the membrane, the NAC domains traverses the alkyl phase as a β-barrel with K80 of the β-turn region interacting with negatively charged lipid head-groups of the trans bilayer while the Nt domains remain on the surface of the cis bilayer as partially-occupied amphipathic α-helices. These types of trimers may serve as precursors for larger transmembrane channels.

### α-Syn transmembrane channels

Lysenin and Aerolysin are the only experimentally-determined protein structures that have concentric β-barrels [31, 32]. Like Lysenin and Aerolysin, α-Syn can exist in a soluble form and interact with membranes to form channels. Atomic force microscopy images of α-Syn channel assemblies [21] are remarkably similar in size and shape to those obtained for Lysenin [66] and Aerolysin channels [67] (Figs. 20 & 23).

**Figure 20.**
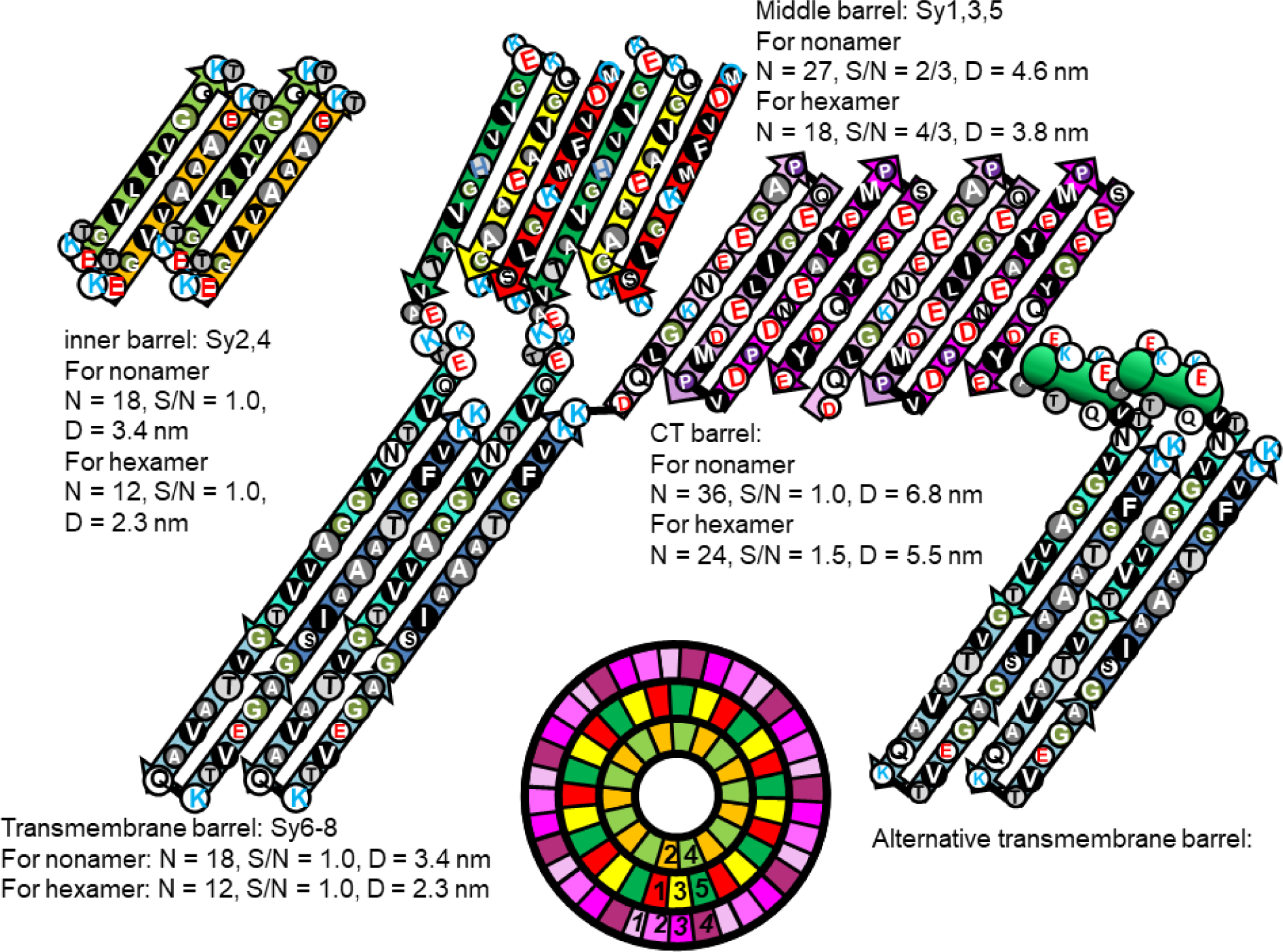
Flattened representations of two monomers for hexamer and nonamer channels. The wedge schematic on the bottom illustrates relative positions of the α-Syn strands in a cross-section of the nonamer’s soluble domain. Segment numbers of the tentative outer Ct β-barrel are italicized. The Nt domain has a Type 1P conformation with the Sy1,3,5 β-barrel in the middle and the Sy2,4 β-barrel at the core. NAC domains form the transmembrane pore; an alternative model of the Sy5 signature region and the NAC pore is illustrated on the right. The NAC β-hairpins are in series with the inner barrel of the soluble domain but are linked to the Sy5 and Ct1 strands of the two outer barrels. Parameters for each barrel are listed.

**Figure 21.**
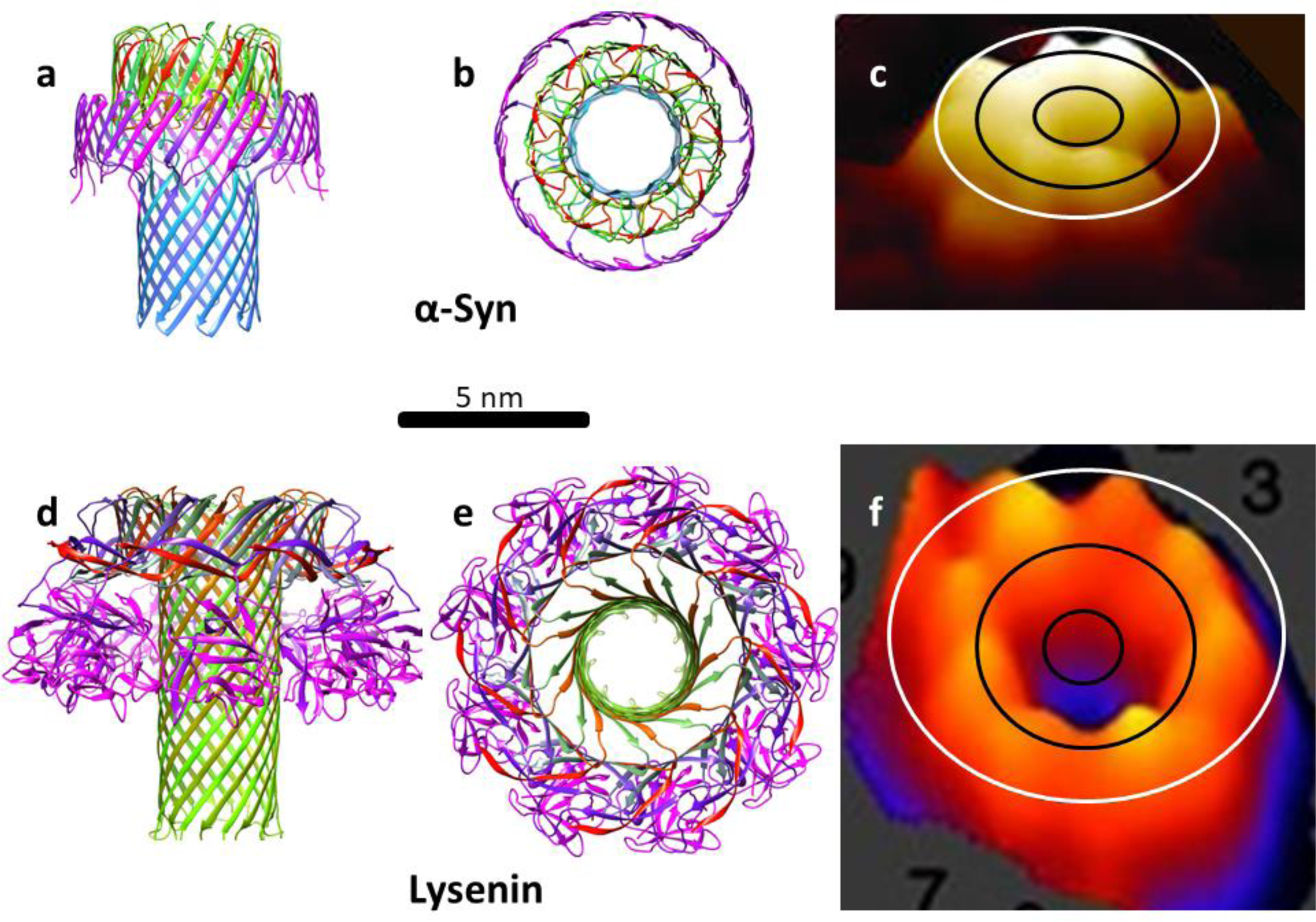
Comparison of our model of a nonamer α-Syn channel to the structure of Lysenin. (a & b) Side and top views of ribbon representations of an atomic scale model of the α-Syn channel. The ribbons are colored by a rainbow spectrum beginning with red for the N-termini and ending with magenta for the C-termini. c) Atomic-force microscopy (AFM) image of the extracellular domain of an α-Syn channel (Adapted from Quist et al. [21]. The ovals represent outlines of the Nt (black) and Ct (white) β-barrel walls of the model. (c & d) Side and top views of a cryo-EM-determined structure of Lysenin [31]. e) An AFM image of the extracellular domains of a Lysenin channel (Adepted with permission from Podobnik M, Savory P, Rojko N, et al [66].

NAC β-hairpins comprise a transmembrane pore and the Nt and Ct domains comprise the soluble domains in our models. However, there are some ambiguities regarding the structures of these domains. The Nt domains could form amphipathic α-helices on the membrane surface or form Type 1P concentric β-barrels as illustrated in Fig. 20. We favor the latter model because it is more like the structures of Lysenin and Aerolysin and appears more consistent with the atomic force microscopy images of α-Syn channels. However, we do not exclude the possibility that the Nt domain forms amphipathic α-helices, possibly as the channels are forming, and that the Ct domain is disordered.

Fig. 20 shows two alternative models for the signature sequence end of Sy5 and the NAC β-hairpin. Both models of the pore resemble the transmembrane region of Lysenin and Aerolysin channels. Although the transmembrane pore-forming regions of Lysenin and Aerolysin are less hydrophobic than the NAC domain of α-Syn and these proteins probably are not strictly homologous, their sequences have some similarities as indicated by the following structural alignment; residues that are identical in two or three sequences are bold, residues with black backgrounds are exposed to lipids in the structures and model, and the underlined regions are the loops connecting the two pore-forming β strands.

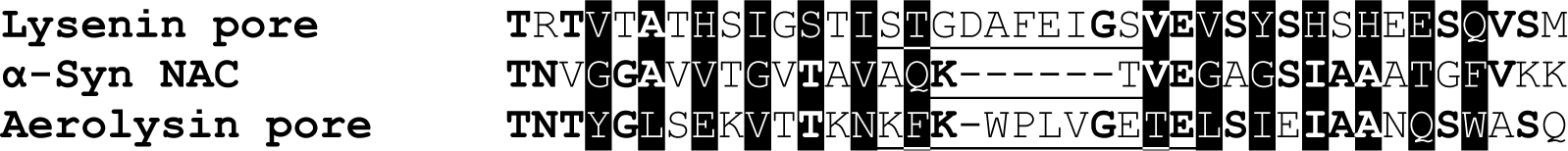

In the first model, the hydrophilic Q79 and K80 side-chains of the putative β-turn within Sy7 would extend into the cytoplasm were the positively charged K80 would likely interact with negative charged lipid head groups. The only other charged residue in this region, E83, extends into the water-filled pore in our model and may increase cation permeation through the pore. In the second model the signature region of Sy5 forms an amphipathic α-helix on the membrane surface and the first putative β-strand (Sy6-Sy7a) is shifted upwardly by two positions. K80 can interact with E83 of the adjacent subunit as well as with lipid head groups. Hydroxyl groups of T72 and S87 extend into the pore in both models. The only polar side-chain groups that would be exposed to lipid alkyl chains are T75 and T92; fewer polar side-chain groups than are exposed to lipids for Lysenin or Aerolysin channels. Although the 11-residue-long β-hairpin of Sy7 is absent in β-Syn, the basic structure could still form with the β-turn occurring at the GVGN residues linking Sy6 to Sy8. This truncation of the putative transmembrane region and removal of positively charged residue at the β-turn may prevent initial interaction with the membrane and/or formation of a channel. If so, this may help explain why β-Syn is not toxic.

### Nonamer channel

Next we developed atomic scale models of the nonamer channel using S/N = 4/3 for the second β-barrel as occurs in the Lysenin structure. Upon examining these models, however, we observed a substantial amount of empty space between the two barrels. Reducing S/N of the middle barrel to 2/3 (the next value permissible for 9-fold symmetry with a three-stranded subunit) reduces the diameter of this barrel by ∼1.2 nm, leaving a gap distance between the walls of the two barrels of only 0.6 nm, the lowest gap distance we allow.

Positioning the outer barrel to reduce clashes among side chains while maintaining the radial symmetry permits H-bonding between the strands of each barrel to be maintained while eliminating empty space between the barrels (Figs. 22 d & f). This tight packing is possible because the majority of side-chains between the barrels are small and the pitches of the pleats are similar enough (7.8 and 7.0 nm) for intermeshing-pleats. The distance between the walls of the middle and tentative outer Ct barrel is greater (∼ 1.2 nm) and the side-chains are larger and more polar. Excluding the last four resides where the conformation is most ambiguous, all charged side chains inside the Ct barrel (D98, K102, E104, D115, E126, E131, D135) form at least one salt bridge with oppositely charged side-chains of the Nt domain.

**Figure 22.**
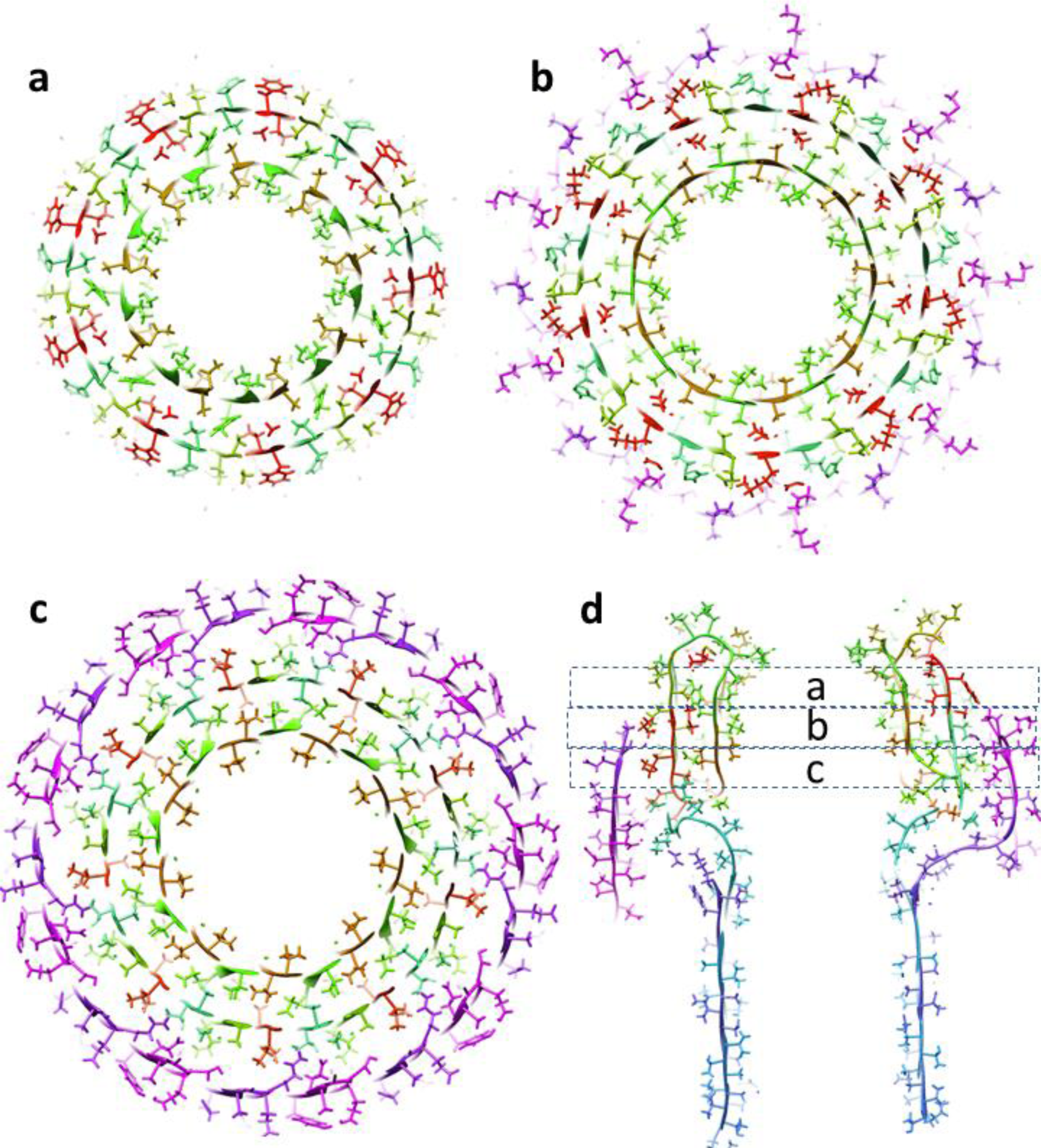
Cross-sections of the α-Syn model’s soluble domain illustrating side-chain packing between the β-barrels. Models are colored as in Fig. 21. (a-c) Three radial cross-sections that are ∼ 0.5 nm apart. (d) Side-view cross-section showing the locations of the radial cross-sections.

**Figure 23.**
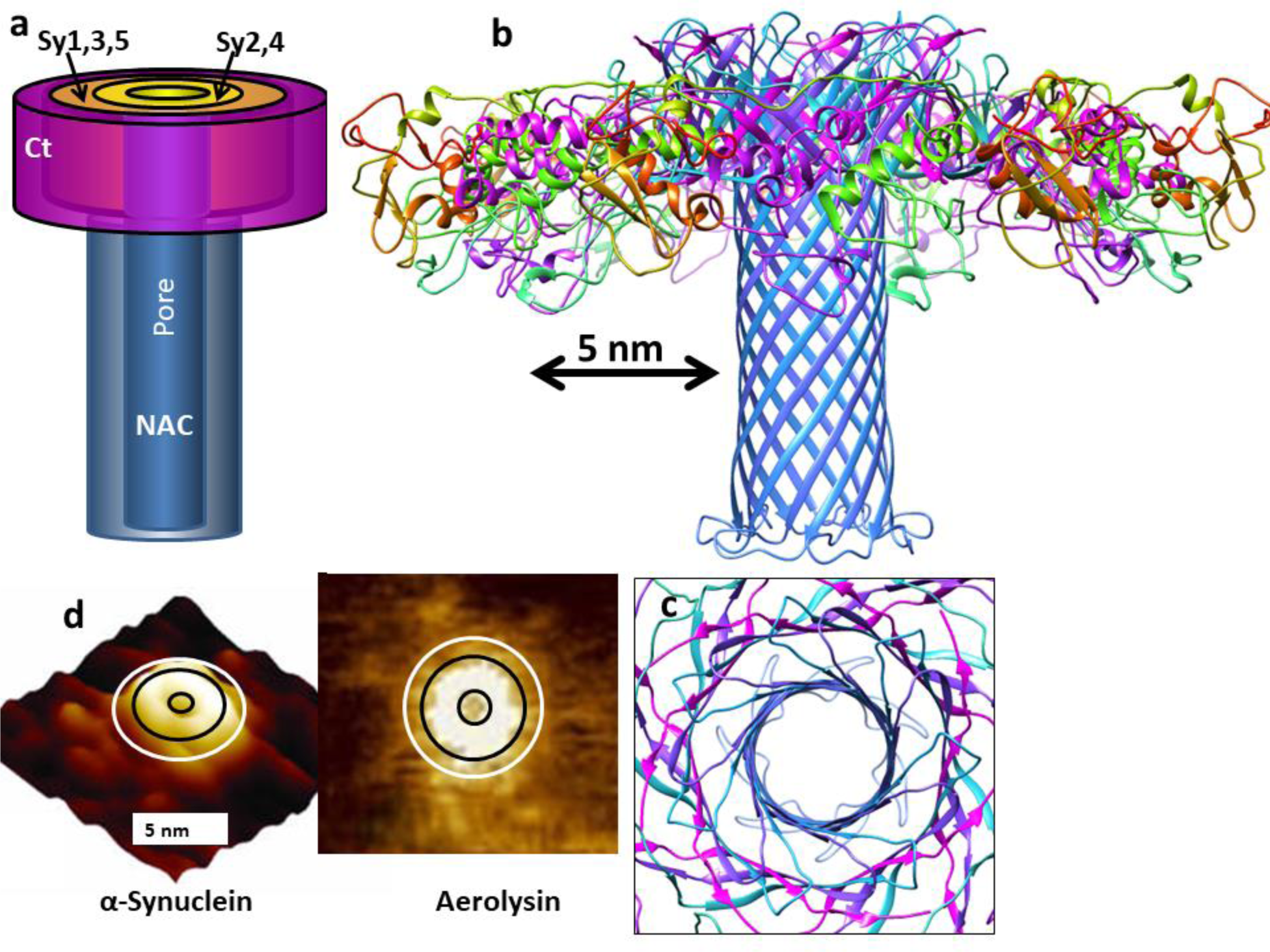
Model of α-Syn hexamer channel compared to the structure of Aerolysin [32]. (a) Schematic of the hexamer channel. (b & c) Side and top views of Cryo EM channel structure of Aerolysin. (d) AFM images of a small α-Syn channel (adapted from Quist et al. [21]) and Aerolysin (Adapted with permission from He J, Wang J, Hu J, et al. [67]). The diameters of the ovals and circles are 6.8, 4.8, and 1.3 nm as predicted for the outer walls and pore of the α-Syn hexamer channel and 7.4, 5.5, and 1.7 nm as calculated for the Aerolysin heptamer.

### Hexamer channel

The basic folding motif of Fig. 20 could also form a hexamer channel similar to that of Aerolysin as illustrate in Fig. 23.

### Cylindrical α-Syn oligomers

Chen et al. [5] obtained cryo-EM images of two toxic cylindrical oligomers of α-Syn.

These oligomers never grow longer. The smaller cylinder has a molecular weight consistent with having 18 monomers. Our model of this assembly resembles that of two channel nonamers stacked end-on with 2-fold perpendicular symmetry (Fig. 24). The primary difference from the channel model is that the NAC β-hairpins from the two nonamers interlock to form a 36-stranded antiparallel β-barrel linking the two nonamers. The dimensions of the model and size of the central pore resemble those of the cryo-EM image.

**Figure 24.**
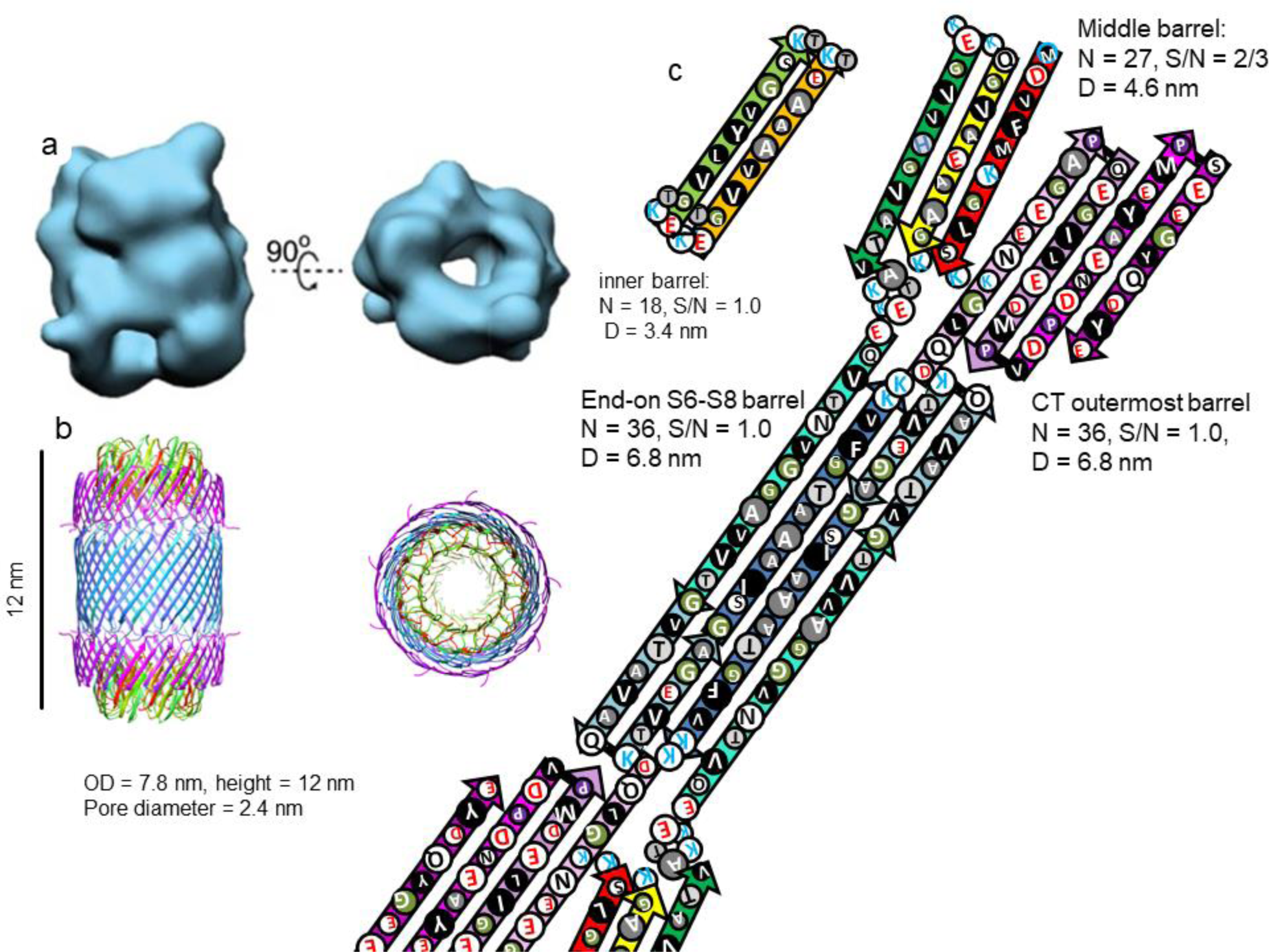
(a) Cryo-EM images of a cylindrical α-Syn 18mer oligomer (copied with permission from Chen et al. [5]) and (b & c) our model of this oligomer. (b) A ribbon representation of our atomic-scale model drawn to the same scale as the Cryo-EM images. (c) Flattened representation of one top and one bottom subunit.

### α-Syn lipoprotein nano particles

α-Syn interacts with negatively charged lipids to form bilayer-like particles with diameters ranging from ∼19 to 28 nm (Fig. 25)[68]. When viewed from the top, the largest of these particles has an ∼6 nm thick ring that probably corresponds to the region occupied by α-Syn. This ring is surrounded by a darker ring, the gray inner ring in turn surrounds a darker circular core that probably corresponds to a lipid bilayer. Experimental findings suggest that the ratio of lipids to protein in some particles is relatively high and that the particles have 8 - 10 α-Syn monomers per particle. CD studies indicates a high degree of α-helical structure. The authors propose that these particles resemble apolipoprotein particles in which amphipathic α-helices reside on the surfaces of the lipid particles.

**Figure 25.**
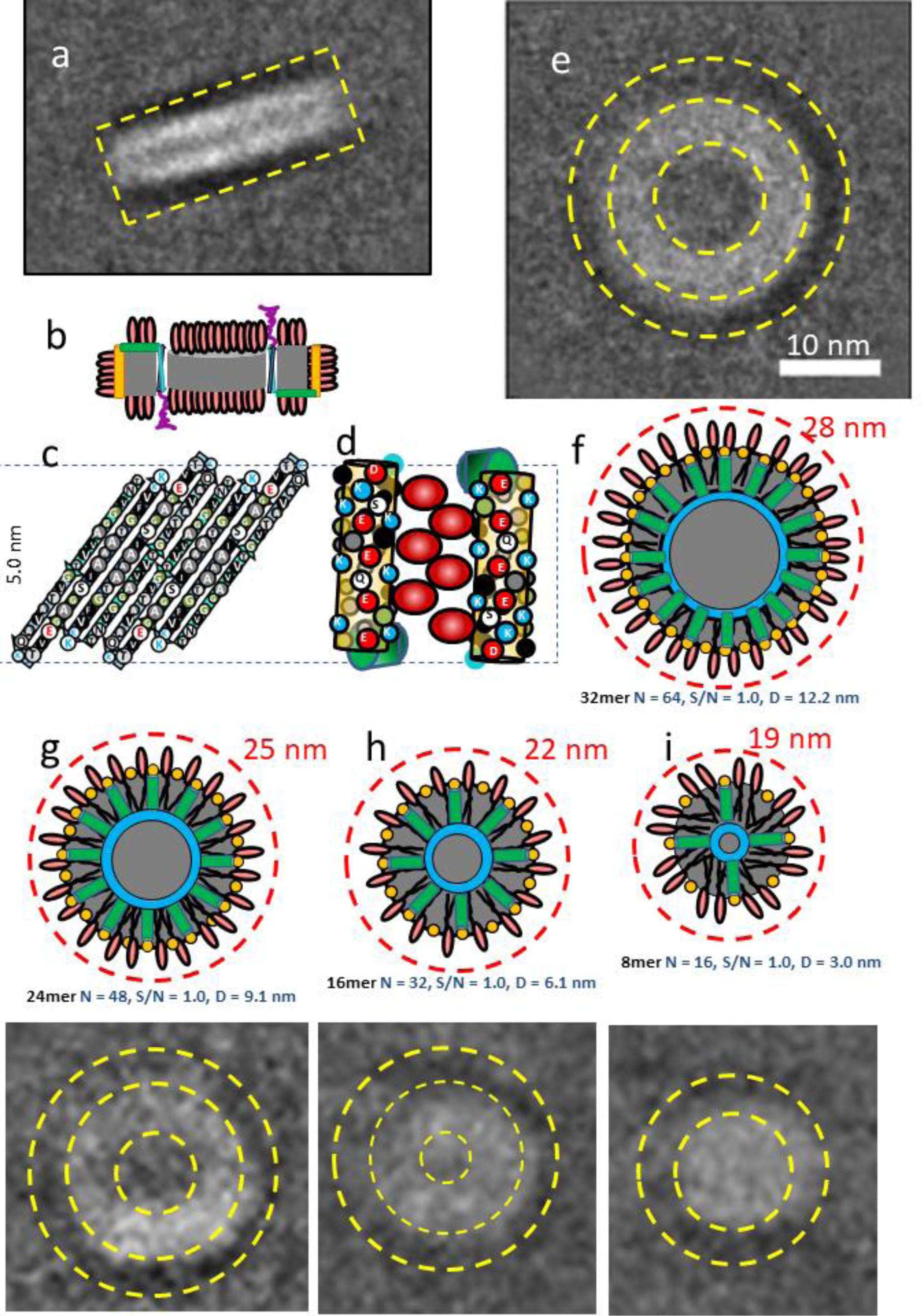
CryoEM images of lipoprotein particles composed of α-Syn and DOPS (1,2-dioleoyl-sn-glycero-3-phospho-L-serine) and αβ-models of these particles. (a) Side view of the largest type of particle showing its bilayer-type configuration. The yellow dashed rectangle shows the dimensions of our model. (b) Schematic of a side view cross-section of our model showing two α-Syn subunits oriented in opposite directions, drawn to the same scale as the EM images. The orange cylinder = Sy1-3 α-helix, green cylinder = Sy3-4 α-helix, cyan and blue arrows = NAC β-hairpin, gray = lipid alkyl chains, red oval = lipid head group. (c) Flattened side view representation of 4 subunits of the NAC barrel. The black dashed lines are 5 nm apart. The only polar side chain atoms in the central region are hydroxyl groups of T72, T75, S87, and T92; only T72 and T92 are on the interior of the β-barrel and T72 can interact with its counterpart at an axis of 2-fold perpendicular symmetry. Other polar side-chains are near the entrances of the barrel and should have energetically favorable interactions; K80 and K96 should interact with lipid head groups, K97 should salt bridge to E83, N65 should H-bond to Q79. (d) Two antiparallel NAC helical regions (colored as in Fig. 5) and their interactions with lipid headgroups (Orange cylinder = Sy1-3 helix, green cylinder = Sy4-5 helix, red ovals = lipid head groups). (e) Top view of a cryoEM image of the largest particle. Yellow dashed circles were added to illustrate the dimensions of the outer circle (D = 29 nm) that surrounds the dark outer ring, and the next two circles (D = 23 & 11 nm) enclose the light ring. (f) Top view of our model of a Type 2 32mer αβ-barrel. Orange circles = Sy1-3 helices, green cylinders = Sy4-5 helices, blue ring = NAC β-barrel (parameters are below image), red ovals = lipid head groups, gray = lipid alkyl chains. (g-i) Similar representations for models and corresponding cryoEM images for 24mer, 16mer, and 8mer. The cryoEM images were copied with modifications from Eichmann et al. [68].

Here we explore a more precisely defined series of models in which the Nt domains form α-helices and the NAC domains form β-hairpins that comprise a Type 2 β-barrel (Fig. 25, b-g). The core of the hydrophobic NAC β-barrel is filled with a lipid bilayer. Sy1-3 segments form amphipathic α-helices oriented approximately parallel to the axis of the β-barrel and positioned about 4 nm from its wall. Lipids fit between antiparallel pairs of Sy1-3 helices where their negatively charged head-groups interact with positively charged lysine side-chains of the helices. Sy4-5 α-helices are approximately perpendicular to the Sy1-3 helices and connect them to the Sy6 β-strand of NAC. Some lipids oriented parallel to those of the central bilayer likely fit between these radial Sy4-5 helices. The smallest of our models has eight α-Syn monomers.

Three larger particles increase in size by steps of eight monomers, consistent with the hypothesis that they form by mergers of smaller particles. The thickness of the putative lipoprotein ring remains constant as the particles increase in size from 8 to 16 to 24 to 32mers. Thus, as the numbers of monomers increase, the increase in the diameter of the assembly equals the increase in the diameter of the β-barrel. If S/N = 1.0 for all β-barrels, an increase of 16 strands (*i.e.,* of 8 NAC β-hairpins) increases the diameter by ∼3.0 nm; *i.e*. if the diameter of the 8mer is 19 nm, then the next three larger particles will have diameters of ∼ 22, 25, and 28 nm. The model dimensions are consistent with the diameters and shapes of the lipoprotein disc CryoEM images.

### Merging of Type 1 oligomers

Fig. 26 shows a hypothetical scheme for how larger Type 1 α-Syn oligomers and channels may develop from Type 1 αβ trimers. In this scheme trimers may merge in solution as illustrated or within the membrane to form hexamers and nonamers, hexamers and nonamers may interact with or within membranes to form transmembrane channels, and nonamers may couple in solution to form octadecamer cylinders. Secondary structure analysis of the cylindrical oligomers [5] indicates the presence of antiparallel β-structure; albeit at a lower percentage than in our model. The dynamic nature of oligomers and the probable simultaneous presence of multiple assemblies and monomers in a prepration of oligomers may complicate evaluation of any specific model based on the results of secondary structure analyses.

**Figure 26.**
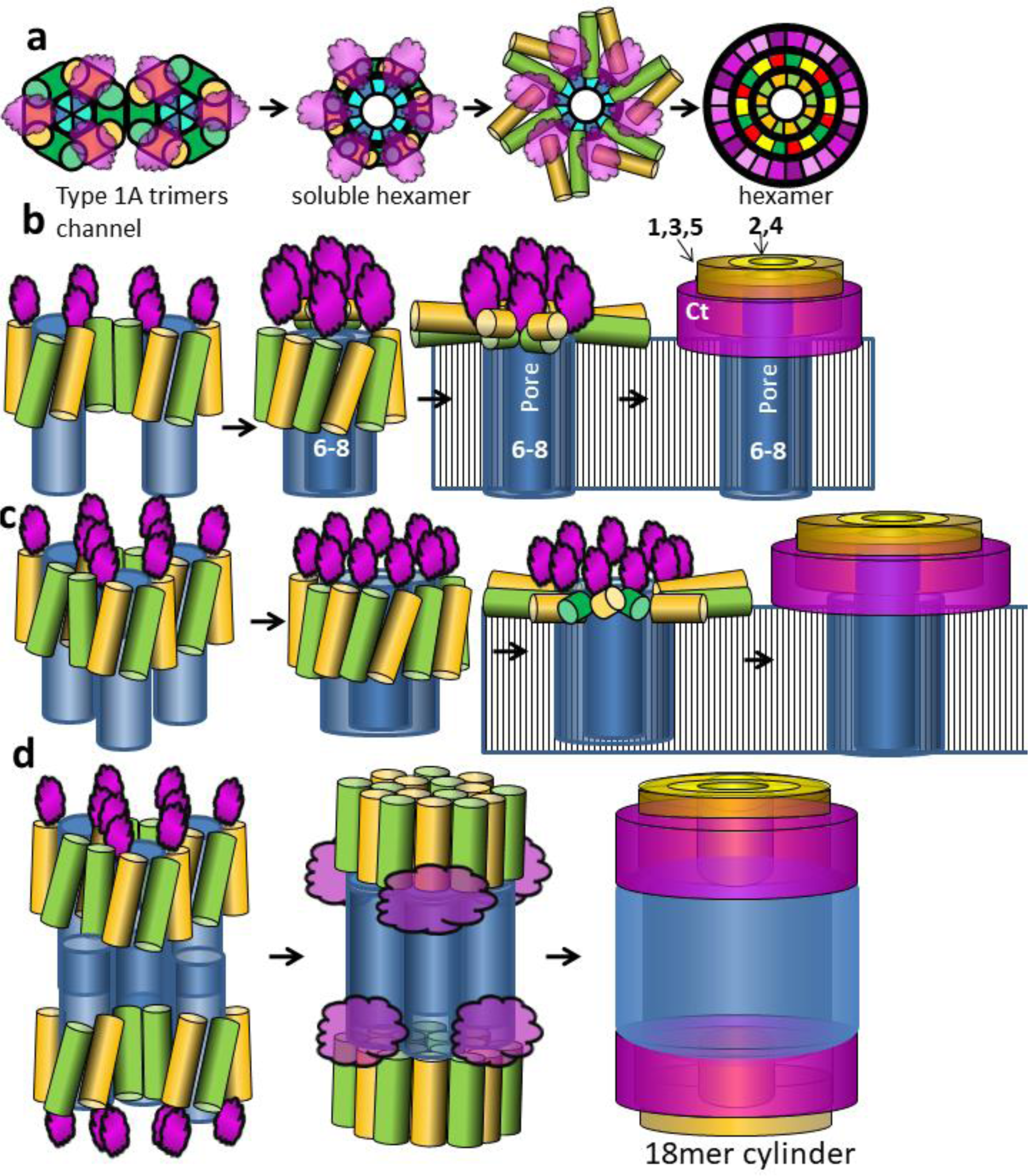
Hypothesis for development of Type 1 oligomers and channels from merging of Type 1 αβ trimers. Small cylinders represent α-helical hairpins of the Nt domain. Larger blue cylinders represent β-barrels formed by NAC domain β-hairpins. Magenta regions represent disordered or β-barrel Ct domains. (a) Wedge representions of assemblies of six α-Syn monomers viewed from the top. Merging of trimers to form hexamers could occur either in solution as illustrated or within the membrane. The NAC domain is not shown in the last schematic. (b) Assemblies of six α-Syn monomers viewed from the side. The black stripped areas represent membrane alkyl regions. When a soluble trimer or hexamer interacts with a membrane, its Nt domain α-helices may peal away form the NAC domain and interact with lipid headgroups on the membrane’s surface, allowing the NAC β-barrel to penetrate the membrane. The Nt and Ct domains may eventually morph into Type 1P β-barrels. (c) Similar assemblies of nine α-Syn monomers. (d) End-on interactins of two soluble nomamers, each formed from three trimers, to form an assembly with 3-fold radial symmetry and 2-fold perpendicular symmetry. Positions of the Nt α-helices, NAC β-barrels, and Ct domain may then shift to more energetically-favorable positions. Next the 6-stranded NAC barrels may merge to form a 36-stranded β-barrel while the Nt helices rearrange to form concentric 18- and 27-stranded β-barrels and the Ct domains may interact to form a partially surrounding 36-stranded β-barrel. The last rearrangement allows all subunits to have identical conformation and gives the 18-mer 9-fold radial symmetry while maintaining the 2-fold perpendicular symmetry.

## Discussion

Our goal has been to analyze whether most experimental results concerning Synuclein oligomers, protofibrils, and transmembrane channels are consistent with relatively simple β-barrel assemblies in which all monomers have approximately identical conformations and interactions. We are as concerned with understanding the overall assembly process as with modeling any specific oligomer structure. Figs. 17 and 25 outline a plausible, if somewhat speculative, series of transitions leading from trimers and tetramers, to larger oligomers, lipoproteins, protofibrils, fibrils, and transmembrane channels.

Although we attempt to make our models and hypotheses as simple as possible, they are more complicated and ambiguous than we would have preferred. Nonetheless, they are no more complex than required to explain experimental results; if anything they are too simple. Our models typically have a higher content of secondary structure than reported experimentally. As previously described for the helical trimers and tetramers of Figs. 5 and 18, we suspect that the secondary structures of these more complex models are also only fractionally occupied. A major point of Figs. 17 and 25 is that the assemblies are dynamic and unlikely to be as structured as the models imply, especially as they merge or morph into other assemblies or break down. Thus we view our models as idealized hypothetical approximations of time-averaged structures for some of the more stable stages of synuclein interactions; i.e. for specific assemblies that can be identified microscopically and/or isolated biochemically.

Tight packing between adjacent β-barrels in our models may account for some features of synuclein sequences that, to our knowledge, have not been explained previously: e.g., the prevalence of small apolar side-chains, especially glycine and alanine, but also valine, and small ambivalent threonine and serine within putative β-strands and the preference for valine over the larger isolucine and leucine side-chains. The high degree of conservation of these residues among the three synuclein families suggests a vital role for these residues, but that role is unlikely to involve fibril formation since β-Syn does not form fibrils. A novel feature of our concentric β-barrel hypothesis is the concept that intermeshing-pleats facilitates tight packing between adjacent concentric β-barrels in some assemblies. The presence of glycines may reduce steric clashes when this type of packing does not occur; i.e., if pleates cross one another. Another striking feature of synuclein sequences is the distribution of charged residues. Although there is evidence that the negatively charged Ct domain is important to some interactions and assemblies, its purpose has not been explained. Here we suggest that its negative charges interact with positively charged components of some zitterionic lipid headgroups, particularly in high-density lipoproteins, but possibly in membranes as well. Highly conserved positively charged lysines of the putative signature turn regions are proposed to interact with negatively charged components of lipids and fatty acids.

The major motivation for analyzing amyloid structures is to develop effective ways to prevent, treat, and/or cure the devastating diseases they cause. Those of us who study molecular structure want to know which assemblies are functional, which are benign, and which are toxic. Then perhaps we can eliminate or neutralize toxic culprits without hindering vital functions. Low resolution EM images and hypothetical models will not suffice; but they may help achieve these goals.

Lysenin and Aerolysin toxins share several properties with the concentric β-barrel models we have proposed previously for Aβ42 and propose here for synuclein amyloid oligomers: 1) they are composed of multiple identical subunits related by radial symmetry, 2) they contain concentric β-barrel structures, 3) diameters and tilt angles of their β-barrels are consistent with β-barrel theory, 4) they exist both in soluble and transmembrane channel conformations, and (5) AFM images of the channel conformations are similar. However, amyloids are substantially more polymorphic and less stable than the two toxins; so much so that they are often said to be intrinsically disordered. These factors complicate experimental determination of their oligomeric structures. In the Supplement we list some examples of experimental approaches suggested by our models. These include experiments to: reduce polymorphism and disorder by stabilizing specific oligomer structures, determine which assemblies are toxic and which portions of those assemblies are responsible for the toxicity, to obtains additional structural images, and to develop techniques and/or drugs to treat PD and other Synuclein-related diseases.

We hope our hypotheses and models will assist efforts to obtain the detailed evidence that can lead to medical breakthroughs.

## Supporting information

experimental tests

## Acknowledgements

This work was supported in part by the Intramural Program of the National Institutes of Health, National Cancer Institute, Center for Cancer Research.

