## Supplementary material for "The amyloid concentric β-barrel hypothesis: models of Synuclein oligomers, annular protofibrils, lipoproteins, and transmembrane channels": experimental tests

### Supplement

#### *Experimental Tests and Targeting Strategies.*

Here we suggest some ways the model could be tested, ways to stablize specific assemblies at the expense of others, and strategies of development of prevention, treatment, and/or cures. We concentrate on testing the broad features and concepts that do not depend upon fine details of the models.

Perhaps the most important model-based hypotheses involve Sy7: 1) that deletion of Sy7 in β-Syn is responsible for most differences between α-Syn and β-Syn, 2) that Sy7 forms a β-hairpin in most oligomers (except hexamers), 3) that the Sy7 β-hairpin is exposed in Type1 oligomers and inserts into the alkyl portions of membranes, 4) that it traverses the membrane in α-Syn channels and forms the trans part of a NAC pore, 5) that it contributes to α-Syn's toxicity, 6) that it is an essential component of the 18mer α-Syn cylindrical oligomers, and 7) that it forms a β-barrel surrounding a central bilayer in some lipoprotein nanoparticles. These hypothese raise the following questions and suggest the following experimental tests:

1) Synthesize an analog of this putative β-hairpin. The β-hairpin structure might need to be stabilized by making the peptide cyclic (near the beginning and end of Sy7) and/or by introducing a disulfide bridge at a region distal from the putative β-turn. Make antibodies to the turn region. Determine whether the antibodies bind to oligomers and if so whether they inhibit the toxicity of α-Syn. Determine whether the Sy7 peptide will bind to β-hairpin-binding antibodies [69,70] and if so use these complexes to determine the structure of the bound Sy7 peptide. Tether Sy7 peptides together with short flexable (e.g. polyglycine) linkers and determine whether in apolar environments they will form 6-, 8-, 12-, or 18-stranded β-barrel structures that can be determined experimentally.

2) Determine whether inserting the Sy7 α-Syn sequence into β-Syn between Sy6 and Sy8 will alter its properties to make it similar to α-Syn; i.e., will the hybrid be toxic, will it insert into membranes to form channels, will it bind fatty acids, will it form fibrils?

3) Determine whether deleting Sy7 from α-Syn makes it behave like β-Syn; i.e., will it no longer be toxic, will it interact with native α-Syn and if so will these mixed oligomers be non-toxic, will it no longer bind fatty acids, will it no longer form fibrils but be able to form oligomers, will it still form high-density lipoprotein stacks of discs?

4) The sequence of the S7 region differs markedly between α-Syn (VT**G**V**T**AV**AQ**K**TVE**G**A**GS) and γ-Syn (FS**G**S**T**NT**AQ**T**TVE**E**A**EN; only 8 (underlined) of 17 residues are identical, it has a zero net charge in α-Syn and a minus three net charge in γ-Syn, and it is more polar in γ-Syn. Will replacing portions of Sy7 of α-Syn with the homologous sequence in γ-Syn reduce its ability to penetrate membranes and/or form transmembrane channels? Will a K80E mutation in the putative Sy7 turn region of α-Syn have a similar effect? If so, will the mutant interact with native α-Syn to reduce its toxcity and can methods such as CRISPR be used to introduce mutations into α-Syn that eliminate toxic effects without affecting vital functions [71]? Will mutating these residues in γ-Syn to those in α-Syn increase its ability to penetrate membranes and/or form channels?

5) **Hypothesis**: NAC domains are proposed to be β-hairpins that assemble to comprise transmembrane β-barrel channels.
1) Determine whether NAC peptides alone will form β-hairpins in aqueous or hydrophobic solvents. If not, will they do so when their N- and C-termini are linked to make them cyclic? Will they form β-barrel channels if multiple peptides are connected with short flexible linkers that should remain in the aqueous phase/lipid headgroup region: e.g., rich in glycine and hydrophilic residues?
2) The pore-forming NAC domain is proposed to have a structure similar to those of Lysenin and Aerolysin channels. The model of the NAC β-hairpin can be matched to the pore-forming transmembrane structure of Aerolysenin and Lysenin to design hybrid proteins. Synthesize hybrid proteins in which the transmembrane region of Lysenin or Aerolysin is replace with the NAC domain and determine whether they will form transmembrane channels. If so, solve the 3-D structure of the hybrid protein.

Synucleins have no native cysteines, raising the possiblity of stabalizing some conformations by introducing disulfide bridges.

1) Type 2 structures with 2-fold vertical symmetry contain residues that are proximal to their counterparts in adjacent subunits at axes of 2-fold symmetry. Thus replacing a residue at one of these positions might covalently link pairs of subunits and stabilize a specific structure while preventing alternative structures. For example, identical residues predicted to by proximal in the Nt domain of Type 2A, Type 2P, and Type 2 αβ-barrel models differ. Differing predictions can also be made for assemblies of the same type but that are composed of differing numbers of subunits or that adopt alternative conformations.

**Hypothesis**: Some hydrophobic side-chains (F4, L8, V26, A30, V48, V52) should be on the exterior of the putative Sy1,3,5 β-barrel formed by Nt in channel and cylinder models (Type 1P). These residues could be important if the Ct domain forms a third β-barrel around the Sy1,3,5 barrel. A Nt peptide would lack the NAC and Ct domains.

1) Will changing these hydrophobic side-chains in a Nt peptide to hydrophilic side-changes stablilize a Type P β-barrel structure similar to those proposed in some channel and cylindrical oligomer models while destabilizing alternative amphipathic α-helical structures?
2) If a specific Type P Nt structure can be stabilized and isolated, can its structure be solved?

**Hypothesis**: The Nt domain has numerous residues that do not favor an amphipathic α-helix conformation.

1) Will mutating these residues to those more favorable for α-helices stabalize an α-β-barrel trimer or tetramer structure for the Nt-NAC domains and/or stabalize lipoprotein nanoparticles; e.g.will mutating G14, G25, G36, G51, T22, T33, T59 of the signature sequences to A and/or mutating A17, Y39, T53 of the polar faces to Q stabalize αβ-barrel structures?

**Hypothesis**: High density lipoprotein assemblies form in the presence of Synucleins and sphingomylein. We suggest that the lipid alkyl chains interact with the hydrophobic interior of the inner Sy2,4 β-barrel for α-Syn and β-Syn, that the lipid's negatively charged phosphate groups interact with lysines of the signature connectors, and that the positively charged moiety of sphingomylein interacts with negative charges on the interior of a Ct β-barrel.

1) Are these assemblies specific for sphingomylein or will they form with most zwitter-ionic lipids?
2) Will mutating some negatively charged residues of the Ct domain to lysine or arginine reduce or eliminate formation of the sphingomylein lipoproteins? If so, which negatively charged groups are more important?
3) The shapes of γ-lipoprotein assemblies differ dramatically from those of α-Syn and β-Syn lipoproteins. We attribute these differencies to major differences in the Ct domain. Will replacing the Ct domain of α-Syn with that of γ-Syn and vice versa change properties of each lipoprotein to resemble the other?
4) Can photo-activated cross-linking groups be added at different locations in sphingolipid analogs? If so, can the synuclein residues and/or segments to which they cross-link be determined?
5) Can the putative antiparallel β-sheet of the Ct domain be stablilized by introducing disulfide bridges at appropriate locations that are predicted to be next to a perpendicular axis of 2-fold symmetry: e.g., K102C, N103C, Y233C, E237C, P238C?
6) Can two lipoprotein assemblies (e.g., two octamers) of adjacent discs be cross-linked by introducing a cysteine at the position nearest the predicted axis of 2-fold symmetry in the Ct domain?

**Hypothesis**: The Ct domain and Sy7 are non-essential components of some Type A models and lipoprotein nanoparticles.

1) Will deleting Ct of α-Syn favor Type A oligomers over Type P oligomers, protofibrils, fibrils, lipoproteins, and channels for which these regions may be important?

2) Will deleting Ct affect formation of lipoprotein nanoparticles?

**Hypothesis**; The N-terminus of the Nt domain is near the C-terminus of the NAC domain in the αβ-barrel trimer model of Fig. 18.

1) Can this structure be stablized by cross-linking the two termini; e.g. by deleting the Ct domain and making the Nt-NAC peptide cyclic, or by using a short flexible connector to link three or four Nt-NAC peptides together?

**Hypothesis**: The last eight residues of the Ct domain are almost identical in α-Syn and β-Syn. These segments from adjacent subunits self-associate in our Type 2P and lipoprotein models.

1) Will deletion or mutations of this segment affect formation of Type2P oligomers, annular and tubular protofibrils, and/or lipoprotein stacks?
2) Can subunits be cross-linked by mutating residues within this segment to Cys? If so, will the cross-links stabilize specific structures?

**Symmetry averaging:** We have found symmetry averaging to be useful in analyzing EM imaging of Aβ42 annular protofibrils (unpublished). If some of the synuclein assemblies described here have the proposed radial symmetry, this approach might improve the resolution of the images.

**Freeze fracture studies:** We have utilized results of preliminary freeze-fracture studies to constrain some of our models of Aβ42 channels. However, these data are limited and to our knowledge this methodology has not been used to analyze other amyloid channel assemblies, including those formed by α-Syn. Such studies might provide additional information about structures of the transmembrane regions of these channels.

**Table S2. Permissible S/N values for symmetric β-barrels used in the models, and approximate values of α and D/N as a function of S/N.**

| **S/N** | **s/uc** | | | | | | | | **α** | **D/N** |
|---|---|---|---|---|---|---|---|---|---|---|
| | **2** | **3** | **4** | **5** | **6** | **8** | **10** | **12** | **(°)** | **(nm)** |
| **0.4** | | | | ■ | | | ■ | | **16** | **0.160** |
| **0.5** | | | ■ | | | ■ | | ■ | **20** | **0.163** |
| **0.6** | | | | | | | ■ | | **23** | **0.168** |
| **2/3** | | ■ | | | ■ | | | ■ | **26** | **0.170** |
| **3/4** | | | | | | ■ | | | **28** | **0.174** |
| **0.8** | | | | ■ | | | ■ | | **30** | **0.178** |
| **1.0** | ■ | | ■ | | ■ | ■ | ■ | ■ | **36** | **0.189** |
| **1.2** | | | | ■ | | | ■ | | **41** | **0.203** |
| **4/3** | | ■ | | | ■ | | | ■ | **44** | **0.213** |
| **1.5** | | | ■ | | | | | | **47** | **0.227** |

For radially symmetric β-barrels the number of permissible values of S/N is a function of the number of strands per radial unit cell (s/uc is the number of strands per subunit for Type 1 structures and twice the number of strands per subunit for Type 2 structures). The black rectangles indicates that the S/N value is permissible. When N ≥ 10, values of the tilt angle, α, and D/N remains relatively constant for any given value of S/N.

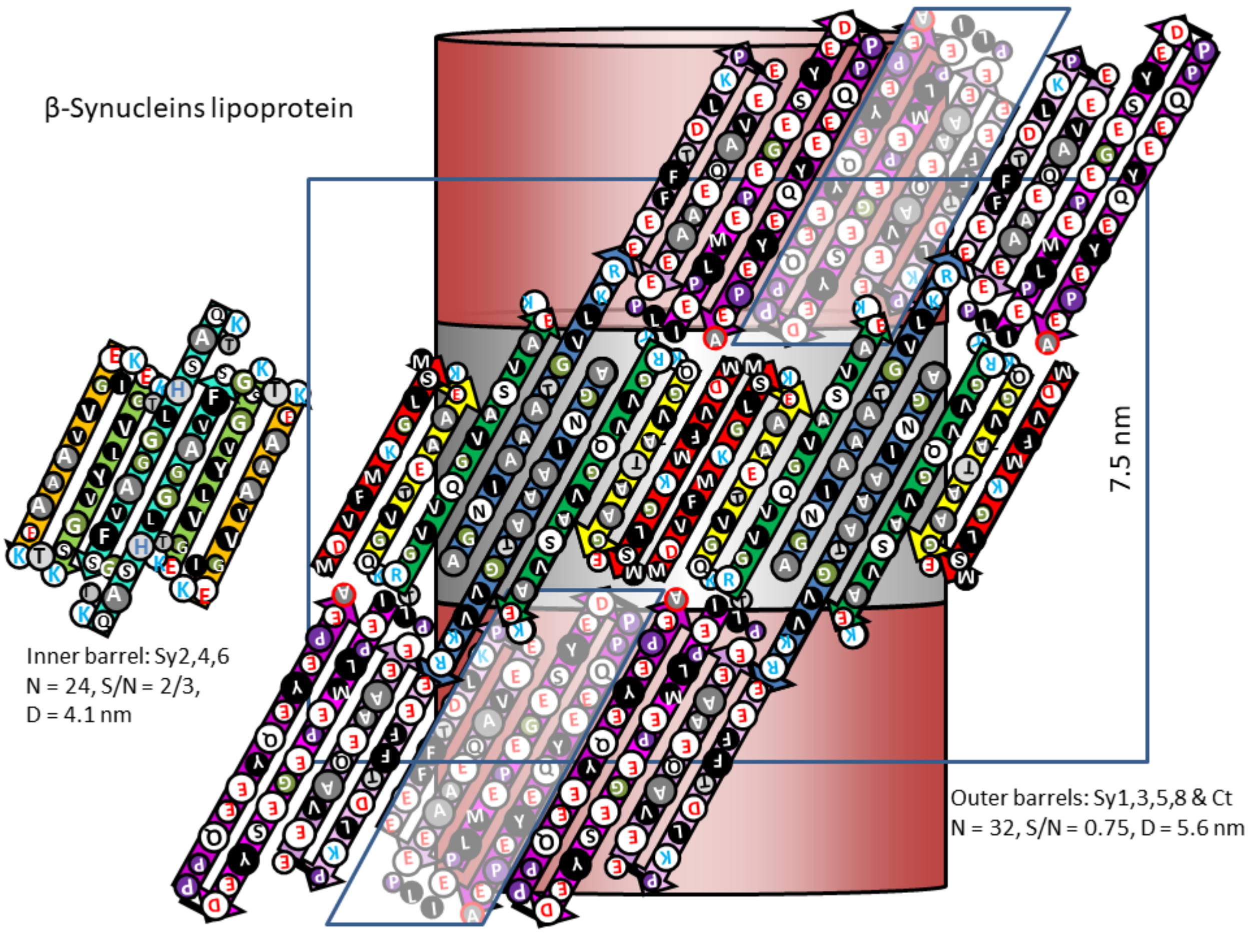


Supplement Figure S1. Schematics of β-Syn stacked discs. Similar to Fig. 15 except the model is for β-Syn.
